# Cracking vacuolar fumarate and malate transport shows its function in Arabidopsis metabolism and growth

**DOI:** 10.64898/2026.03.30.714522

**Authors:** Roxane Doireau, Elsa Demes-Causse, Paloma Cubero-Font, Solenne Berdarocco, Younès Dellero, Alexis De Angeli

## Abstract

Malate and fumarate constitute a significant transient carbon stock that is dynamically synthesized during the photoperiod. These organic acids are diurnally stored and remobilised from the vacuole, and they have a key role in the cellular metabolic regulation. This function is well known in C4 and CAM plants. However, in C3 species that are the majority of terrestrial plants, the importance of the vacuolar accumulation/release and its influence on plant growth is still an open question. In Here we addressed this issue generating multiple knockout mutants in *Arabidopsis thaliana* lacking vacuolar anion channels of the Aluminium-Activated Malate Transporter (ALMT) family, to impair malate and fumarate transport to the vacuole. We show that in these mutants reducing vacuolar transport of malate and fumarate in mesophyll cells leads to a dramatic growth impairment. Metabolic and fluxomic analysis revealed that vacuolar malate and fumarate transport influences plant carbon and nitrogen metabolism as well as cellular pH and ionic homeostasis. In conclusion, our results show that the transport organic acids like malate and fumarate across the vacuolar membrane is essential for plant growth in a C3 plant too. These results establish the importance of the vacuolar pools of malate and fumarate in plant metabolism.

## INTRODUCTION

Land plants store a significant part of the carbon (C) fixed by photosynthesis as starch, sugars and organic acids (Gibon *et al*., 2009). The main organic acids produced and accumulated are malate^2-^, citrate^3-^ and, in some plants including Arabidopsis, fumarate^2-^ (Chia *et al*., 2000). These three organic acids are key metabolites at the crossroad of several metabolic pathways (Britto & Kronzucker, 2005; Hurth *et al*., 2005; Zhang & Fernie, 2023). The importance of malate synthesis and accumulation is well known in C4 and CAM (Crassulacean Acid Metabolism) plants, where it is the keystone metabolite for these specialised metabolisms adapted to warm climates (Winter & Smith, 2022). However, most of the land plants are C3. They produce/accumulate malate^2-^ and fumarate^2-^ during daytime (Gibon *et al*., 2009) but, compared to C4 and CAM, they accumulate much less malate^2-^ and fumarate^2-^ (Winter & Smith, 2022). Further, while carbon allocation and plant growth were shown to be coordinated (Sulpice *et al*., 2014), the importance of organic acid vacuolar accumulation in C3 plants growth is unclear.

Within the cell, malate^2-^ and fumarate^2-^ are synthesised in the mitochondria by the TCA cycle and, in the cytosol, malate^2-^ is produced by the PEP carboxylase (PEPc) in anaplerotic CO2 fixation (Ludwig *et al*., 2024). In *Arabidopsis thaliana*, which is a C3 plant, a significant part of the malate^2-^ produced during daytime is reduced to fumarate^2-^ in the cytosol (Pracharoenwattana *et al*., 2010). Indeed, in Arabidopsis a cytosolic FUMARASE2 (FUM2) converts malate^2-^ to fumarate^2-^ (Pracharoenwattana *et al*., 2010). Both, malate^2-^ and fumarate^2-^ form a significant stock of carbon fixed during the day and used at night (Araújo *et al*., 2011). These two organic acids are involved in several metabolic pathways and in the pH regulation of different intracellular compartments (Britto & Kronzucker, 2005; Hurth *et al*., 2005; Zhang & Fernie, 2023). Further, malate^2-^ and fumarate^2-^ participate to the turgor pressure regulation (Rentsch & Martinoia, 1991). The multiple functions of malate^2-^ require a tight control of its cytosolic concentrations, and an important process to regulate cytosolic malate^2-^ and fumarate^2-^ homeostasis is through their accumulation into the vacuole (Martinoia *et al*., 2007). Further, cytosolic fumarate^2-^ synthesis in the cytosol and accumulation in the vacuole is proposed to be a metabolic fail-safe to keep cytosolic malate^2-^ homeostasis (Saunders *et al*., 2022).

In the leaves, the vacuole of the mesophyll cells is the subcellular compartment storing the major part of these organic during the day (Destailleur *et al*., 2021). Thus, the vacuolar transporters mediating malate^2-^ and fumarate^2-^ fluxes play a key function for the cytosolic homeostasis of these organic acids. There are only a few malate^2-^ and fumarate^2-^ transporters identified in the vacuolar membrane, the tonoplast Dicarboxylate Transporter (tDT) and four members of the ALMT (Aluminium Activated Malate Transporter) family (Kovermann *et al*., 2007; Meyer *et al*., 2011; Eisenach *et al*., 2017; Medeiros *et al*., 2017; Doireau *et al*., 2024). So far, it was shown that knocking out *AttDT* or *AtALMT5* affects the total content of malate^2-^ or fumarate^2-^ in Arabidopsis. Indeed, the *AttDT* and *AtALMT5* knock-out display reduced malate^2-^ and fumarate^2-^ storage, respectively (Medeiros *et al*., 2017; Doireau *et al*., 2024). However, despite this accumulation defect none of these two mutants shows impaired growth. Notably, plants knocked-out for the other vacuolar *At*ALMTs (*AtALMT4* and *AtALMT9*), which are expressed in the mesophyll, do not show significant reduction of organic acid accumulation or plant growth (Baetz *et al*., 2016; Eisenach *et al*., 2017). Therefore, the importance of malate^2-^ and fumarate^2-^ vacuolar accumulation in Arabidopsis and other C3 plants is still an open question.

In the present work, we tackle the question of the importance of vacuolar malate^2-^ and fumarate^2-^ transport and accumulation in the C3 plant *Arabidopsis thaliana* and its role in the metabolism. We generated multiple knock-out mutants of the vacuolar *At*ALMT channels, focusing on those expressed in the leaves and in the mesophyll, *At*ALMT4, *At*ALMT5 and *At*ALMT9 (Kovermann *et al*., 2007; Eisenach *et al*., 2017; Doireau *et al*., 2024). We found that *almt5almt9* and *almt4almt9almt5* knock-out plants are strongly impaired in their growth. These defects were associated with a dramatic reduction of fumarate^2-^ and malate^2-^ content in the shoots. The capacity to transport fumarate^2-^ across the tonoplast was dramatically reduced. The loss of the three vacuolar *At*ALMTs resulted in a reduced potassium content and is associated to a more alkaline cell-sap pH. Metabolomics analysis revealed an impact on the carbon and nitrogen metabolism in *almt5almt9* and *almt4almt9almt5* mutants. Our data show that in the mesophyll, the vacuolar *At*ALMT are the main regulators of malate^2-^ and fumarate^2-^ accumulation during daytime. Further, these data demonstrate that the transport of fumarate^2-^ across the vacuolar membrane participate to the regulation of the plant cell central metabolism in C3 plants.

## MATERIALS AND METHODS

### Plant material and growth conditions

*Arabidopsis thaliana* genotypes were: Columbia-0 (Col-0), *almt5-1* (WiscDSLox386E04.0), *almt9-1* (SALK_055490), *almt4* (SALK_086236), and *almt5almt9*, *almt4almt5*, *almt4almt9*, *almt4almt9almt5* obtained by crossing the previous genotypes. *A. thaliana* plants were grown in pots in a growth chamber with a day/night regime of 8h/16h, 25°C/20°C, 65% humidity and LED lights 150 µmol.m^-2^.s^-1^. Seeds were stratified 72h at 4°C in the dark. For seed propagation, plants were grown in a greenhouse.

### Arabidopsis crossing

The two knock-out alleles *almt5-1* (WiscDSLox386E04.0) and *almt9-1* (SALK_055490) were used to generate the double mutant *almt5almt9*. The double mutant *almt4almt9* was generated using the two null alleles *almt4* (SALK_086236) and *almt9-1*. The double mutant *almt4almt5* was generated using the two null alleles *almt4* and *almt5-1*. The triple mutant *almt4almt9almt5* was generated crossing the double mutants *almt4almt5* and *almt4almt9*. All lines were genotyped by Polymerase Chain Reaction (PCR) and gene expression was analysed by RT-qPCR.

### Screening of T-DNA lines by PCR

T-DNA insertion lines were screened by PCR to confirm the presence of the T-DNA and to verify if the lines were homozygous for all the T-DNA insertion. For this, DNA extraction from leaves was performed with the REDExtract-N-Amp Plant PCR kit (Sigma-Aldrich, Co., MO, USA) followed by PCR with GoTaq® DNA Polymerase (Promega, USA) and specific primers (Supporting Table 3).

### RNA extraction and gene expression analysis

Leaves number 9 of 40 days-old plants were collected and frozen in liquid nitrogen in 2 ml tubes containing two steel beads (2.5 mm diameter). Tissues were grinded for 45 s at 35 s-1 in a mixer (MM400, Retsch, Germany). RNA was extracted from leaves using TRIzol reagent (MRC, USA). Subsequently, 1 µg of RNA was used to perform reverse transcription in the presence of Moloney murine leukemia virus reverse transcriptase (Invitrogen, USA), following the manufacturer’s instructions. Annealing was done with an Oligo(dT)20 primer. The quality of cDNA was verified by PCR using specific primers spanning an intron in the gene ACT2 (At3g18780.2) with the following primers: forward 5’-TGGAATCCACGAGACAACCTA-3’; reverse 5’-TTCTGTGAACGATTCTGGAC-3’. Gene expression was determined by quantitative real-time PCR (qRT-PCR; LightCycler 480, Roche Diagnostics, Switzerland) using TB Green Premix Ex Taq (Tli RNaseH Plus; TaKaRa, Japan) according to manufacturer’s instructions with 2µl cDNA in a total volume of 10µl. All expression values were standardised to the reference gene, YLS8 (At5g08290) (Czechowski *et al*., 2005). The normalised expression was calculated according to (Taylor *et al*., 2019). Gene specific primer sequences are in Supporting Table 3.

### Anion content measurement by High Pressure Chromatography (HPIC)

Entire rosettes of 10 plants of 40 days-old were collected at the end of the photoperiod (ED) in the growth chamber. The rosettes were frozen at -20°C for 20 min, heated at 70°C for 30 min with 5 to 20 ml of H2O (dependant of the size of the rosette) and centrifuged at 18000 g at 4°C for 15 min. The supernatant collected was analysed in a 2 mL vial kit Dionex® (Thermofisher Scientific, USA). Analysis was done by HPIC (Dionex™ ICS-5000+ Capillary HPIC™) using the software Chromeleon (Thermofisher).

### Measurement of Cell Sap pH

Cell sap was extracted from rosette leaves of 5-week-old plants grown in short days conditions (8h light). The entire rosette was chopped off and squeezed in a 2 ml filtered Microtube with a micropestle. Samples were centrifuged for 3 min at 14.000 rpm. The pH of the extracted solution was immediately measured using a pHmeter microelectrode (HI1083, Hanna Instruments).

### Cations and Metal ion Quantification

For metal ion measurements, samples were harvested and dried at 65°C for one week before grinding with a mortar and pestle. Between 5 and 20 mg of samples are homogenised with 750 μl of nitric oxide (65% [v/v]) and 250 μl of hydrogen peroxide (30% [v/v]). Following an overnight incubation at room temperature, the samples were incubated at 85°C in HotBlock (Environmental Express) for 12 to 24 hours. Samples were then diluted by adding 4 ml Milli-Q H2O before the measurement. The following 10 elements were quantified: calcium (Ca), copper (Cu), iron (Fe), potassium (K), magnesium (Mg), manganese (Mn), sodium (Na), phosphorus (P), sulphur (S) and zinc (Zn). Analysis of ion content was performed using ICP- OES (inductively coupled plasma optical emission spectrometer, ICP OES 5800, Agilent Technologies).

### Carbon and Nitrogen quantification by mass spectrometry

Entire rosettes of 7-25 plants were collected 5h after the beginning of the light period in a growth chamber, in short day conditions (8h light). Samples were analysed by isotope mass spectrometry (Vario-PYROcube Elemental Analyser coupled to an IsoPrime Precision mass spectrometer, Elementar, UK). Dried and ground samples were placed in a tin cup and injected into an elemental analyser (Vario-PYROcube, Elementar, UK). After combustion at 920°C in the presence of oxygen and CuO, gas molecules from the sample (mainly H2O, CO2, N2 and Nox) are transported by a carrier gas stream (Helium ultrapur, AirLiquide) to a reduction furnace where nitrogen oxides are reduced to N2 in the presence of copper at 600°C (Dumas reaction). The H2O produced is trapped by sicapent columns (Merck). N2 and CO2 are then separated. The CO2 is trapped at room temperature in a TPD (Temperature Programmable Desorption) column, then released at 100°C. A thermal conductivity detector (TCD) is used to quantify total N and C. N2 and CO2 are then analysed by mass spectrometer (IsoPrime Precision, Elementar, UK) to determine ^15^N and ^13^C isotopic abundances.

### 13CO2 labelling and GC-MS analyses

The experimentation was done using 5–6-week-old *Arabidopsis thaliana* plants, grown in pots in short day conditions (8h light). Entire rosettes were collected before or after 4h of ^13^CO2 incorporation (n = 4 rosettes /genotype/condition). ^13^CO2 labelling experiment was achieved in a custom gas-exchange chamber connected to the LI 6800-F using the 6800-19 Custom Chamber Adapter (LiCOR). The custom chamber had a volume of 3.1 dm^3^, contained 2 small fans that allowed a 95% air renewal in approximately 5 min (Fig. S1) and was placed under a light source (200 μmol photons.m^−2^.s^−1^). The structure of the custom chamber allowed to isolate the rosette leaves during the labelling, without altering plant physiology. Plants were acclimated for 1 hour inside the chamber with ^12^CO2 prior to start the labelling with ^13^CO2 (99% ^13^C, Eurisotop). Standard environmental parameters were controlled by the LI 6800-F, as follows: a leaf temperature of 22-24◦C, 65–80% relative humidity (leaf VPD ranging from 0.8 to 1.2), 430 μL CO2.L^−1^, 0.21 L O2.L^−1^ and a flow rate of 30 cm^3^.s^-1^. Samples were harvested in the light using liquid nitrogen spraying. Rosette leaves were then freeze-dried and grinded into a fine powder. For each sample, 2 mg were used for absolute quantification of net ^13^C fixed using EA-IRMS, and 10 mg for the analysis of metabolite content and ^13^C-enrichment by GC-MS. Polar metabolites were extracted with a mixture of chloroform/methanol/water containing Norleucine and Adonitol as internal standards. Sugars were analysed as TMS- derivatives and organic and amino acids as TBDMS-derivatives following the procedures described in (Dellero *et al*., 2024; Aubert *et al*., 2025). Metabolites are denoted according to the carbon backbone harboured by the fragment analysed (C1C5 means carbons 1,2,3,4,5 for example). Raw data were corrected for natural isotope abundance using IsoCor and dedicated files (Millard *et al*., 2019; Dellero and Filangi, 2021; Le Féon *et al*., 2023). Quantification was achieved with external standard curves of authentic standards and by summing all isotopologues. ^13^C allocation was expressed in µmol^13^C.g^-1^DW.h^-1^ following calculations described in (Aubert *et al*., 2025).

### Protoplast and vacuole isolation

Arabidopsis mesophyll protoplasts were isolated enzymatically from 4-5 weeks old plants. After 60 min incubation in enzyme solution (0.5% (w/v) cellulase R-10, 0.05% (w/v) macerozyme R-10, 1 mM CaCl2, 10 mM MES, pH 5.3 adjusted with KOH, π = 550 mOsm adjusted with D-sorbitol) cells were washed twice and resuspended with a washing buffer (1 mM CaCl2, 10mM MES, pH 5.3 adjusted with KOH, π = 550 mOsm adjusted with D-sorbitol). Protoplasts were kept on ice until used for patch-clamp experiments and vacuoles were released by osmotic shock.

### Patch-clamp recordings

Currents were recorded using an EPC-10 amplifier (HEKA Electronics,Lambrecht/Pflaz, Germany). Data were acquired and analysed using PATCHMASTER and FITMASTER softwares (HEKA). Microelectrodes (resistance 2.5–5 MΩ) were prepared from GC150TF-15 glass capillaries (Harvard Apparatus, Holliston, MA, USA), treated with sigmacote (Sigma-Aldrich Co.) and coated with Sylgard (WPI, Friedberg, Germany). The reference electrode was placed in 1 M KCl agar bridge with 2% agar. Vacuolar side buffer: 11.2 mM DL-malic acid, 100 mM HCl, pH 6 with BisTrisPropane (BTP), p = 550 mOsmol kg^-1^. The cytosolic side buffers were composed of (1) 100 mM Fumaric acid, 0.1 mM CaCl2 (2) 100 mM DL-malic acid, 0.1 mM CaCl2.

For both cytosolic side buffers, pH 7.5 was adjusted with BTP and p = 500 mOsmol kg^-1^ with D-sorbitol. The cytosolic side buffers were exchanged using a gravity-driven perfusion system. Ion currents were recorded in the whole-vacuole configuration. Vacuoles were isolated by osmotic choc from mesophyll protoplasts. Measurements were done ≥ 15 min after the establishment of the whole-vacuole configuration to ensure dialysis of the vacuolar lumen. In our recording conditions, the liquid junction potential was higher than 2 mV, it was corrected accordingly for both tested cytosolic conditions. In both cytosolic conditions, currents were recorded using a voltage-pulse protocol starting at a holding voltage of 0 mV, 2 s lasting voltage pulses were applied in 20mV increments followed by a postpulse for 500 ms and back to 0 mV. In fumarate condition (1), voltage pulses were applied from +35 to -125 mV followed by a postpulse at +35 mV. In the malate condition (2), voltage pulses were applied from +33 to -127 mV followed by a postpulse at +33 mV. Currents were recorded at a sampling rate of 100 µs and low-pass filtered at 1 kHz. Patch-clamp data are presented according to the endomembrane convention for electrical measurements (Bertl *et al*., 1992).

## RESULTS

### *At*ALMT channels are essential for plant growth

*At*ALMT4, *At*ALMT5 and *At*ALMT9 are the three main anion channels present in the vacuolar membrane that transport malate^2-^ and fumarate^2-^ across the tonoplast. They display overlapping expression patterns, specifically in the leaves where they are all expressed in the same tissues (Kovermann *et al*., 2007; Eisenach *et al*., 2017; Doireau *et al*., 2024). To understand the role of the accumulation of these organic acid in leaves we generated a combination of double mutants and a triple knock-out mutants for *AtALMT4*, *AtALMT5* and *AtALMT9* (Fig. 1A; Fig. S2A-B). Knock-out of each gene was confirmed by genotyping and the extinction of gene expression (Fig. 1E-G). The *almt5almt9* and *almt4almt9almt5* plants displayed a drastic reduction of rosette size and biomass and a yellowing of the oldest leaves (Fig. 1A-D and Fig. S2C). Compared to the wild-type, the FW of *almt5almt9* and *almt4almt9almt5* mutants decreased by 84% and 97% respectively (Fig. 1B). Complementation of the double mutant *almt5almt9* with either ALMT5 or ALMT9 expression almost completely rescued this phenotype (Fig. S3A-E). Differently, the *almt4almt9* and *almt4almt5* mutants displayed only a moderate FW reduction of 27% and 37%, respectively. The DW followed the same trend (Fig. 1C) and the FW/DW ratio was reduced in all the multiple mutants but with a stronger decrease in *almt5almt9* and *almt4almt9almt5* (Fig. 1D). I*n vitro* assays showed that in all the multiple vacuolar ALMT knock-outs, the root growth was not affected (Fig. S3F-G).

**Figure 1:**
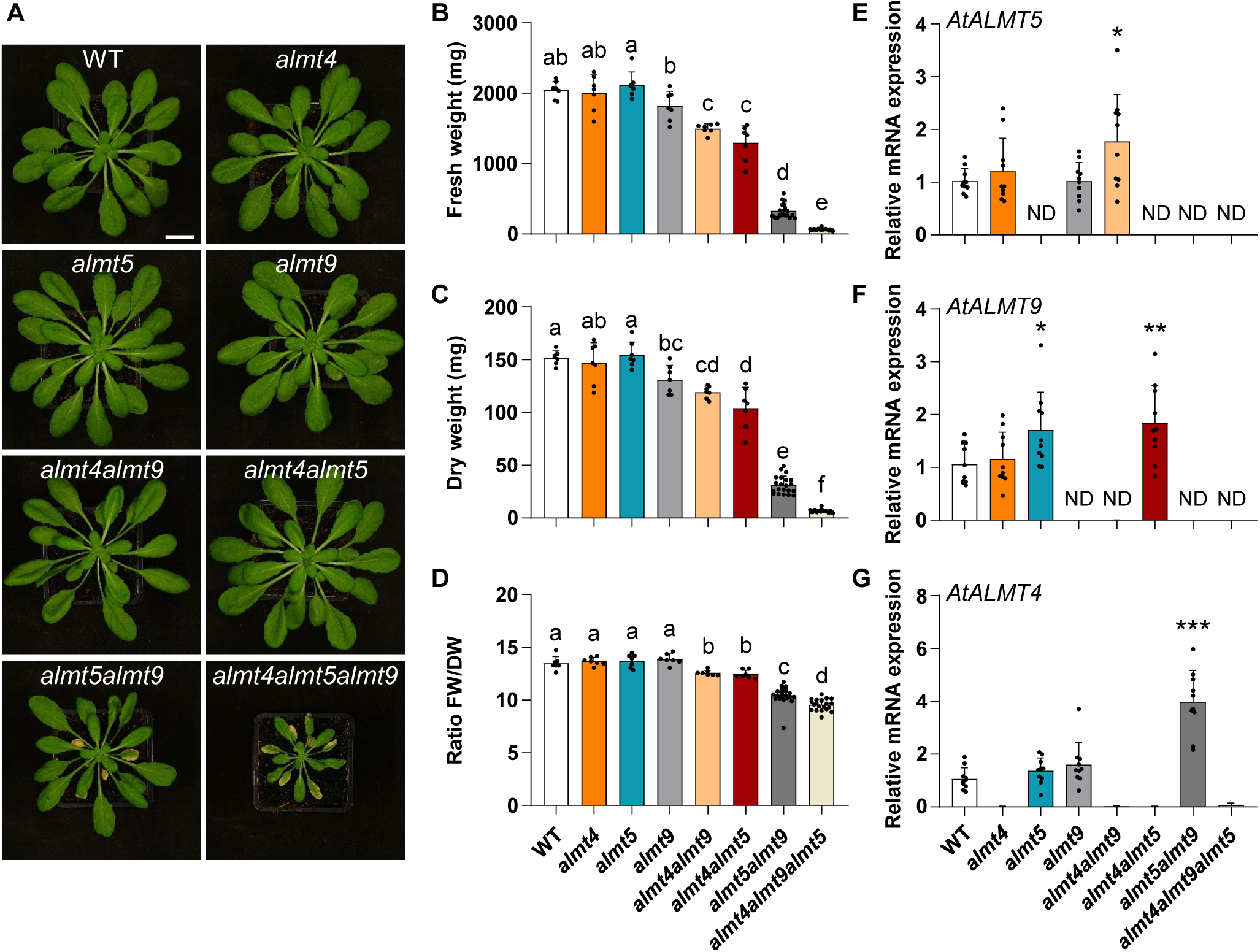
*At*ALMT channels are involved in plant growth. (**A**), *Arabidopsis thaliana* wild-type (WT), simple mutants *almt4*, *almt5*, *almt9*, double mutants *almt4almt9*, *almt4almt5* and triple mutant *almt4almt9almt5* 39 days-old plants grown in short days conditions (8h light). Bar = 2cm. (**B-D**) Rosette fresh weight (FW) and dry weight (**C**) and ration of FW/DW (**D**) of *Arabidopsis thaliana* 40-days old plants knock-out lines grown in short days conditions (8h light). Values are mean + SD, n= 7-21 plants for each genotype. Letters indicate a significant difference (P<0.05). Statistical analysis : Kruskal-Wallis test + Mann-Whitney post-hoc test, Benjamini-Hochberg correction. (E-G) *AtALMT5* (E), *AtALMT9* (F) and *AtALMT4* (**G**) expression quantified by qRT-PCR. 40 days-old plants were grown in short days conditions (8h light). Leaves number 9 were collected at the end of the day (ED). For each genotype 10 plants were collected and analysed pooled by two. Transcript accumulation was quantified after normalisation by the reference gene Yellow leaf specific gene (YLS8). Statistical analysis: Mann-Whitney test (*P<0.05; **P<0.1). ND: non detected.

Our phenotypic analysis suggested a certain degree of redundancy between these three ALMT channels, being more prominent between *At*ALMT5 and *At*ALMT9. RT-qPCR analysis of the wild-type plants revealed that *AtALMT9* and *AtALMT5* were 4 times and 53 times more expressed than *AtALMT4,* respectively (Fig. S3H), which correlates with the phenotypic analysis of the double and triple mutants. Compared to the wild-type, in *almt4almt9* the expression of *AtALMT5* was 1.74-fold higher, while in *almt4almt5* the expression of *AtALMT9* was 1.73-fold higher (Fig. 1E-F).

Interestingly, in *almt5almt9* leaves the expression of *AtALMT4* was 2.74 times upregulated compared to the wild-type (Fig.1G), however this higher expression was not sufficient to recover the growth defect of this mutant. Notably, in non-senescent leaves from *almt5almt9* and *almt4almt9almt5* the expression of the early senescence marker *AtSAG13* was 13 and 119 time higher that in wild-type, respectively (Fig. S2E). Differently, the expression of *AtRBCS1A,* a marker of photosynthetic deficiency (Bresson *et al*., 2018), was not significantly modified between the genotypes (Fig. S2D).

### Vacuolar fumarate transport capacity and storage are affected in multiple knock-out mutants

*At*ALMT9, *At*ALMT5 and *At*ALMT4 are ions channels mediating malate^2-^ and fumarate transport (Kovermann *et al*., 2007; De Angeli *et al*., 2013a; Eisenach *et al*., 2017). Interestingly, *in vivo At*ALMT5 is involved only in fumarate^2-^ accumulation, and patch-clamp analysis of *almt5* mesophyll vacuoles revealed a reduced fumarate^2-^ current density compared to wild-type (Doireau *et al*., 2024; Fig. 2C and Fig. S4A-B). We therefore used whole-vacuole patch-clamp to investigate fumarate^2-^ and malate^2-^ currents in the single and multiple ALMT mutants (Fig. 2 and Fig. S4).

**Figure 2:**
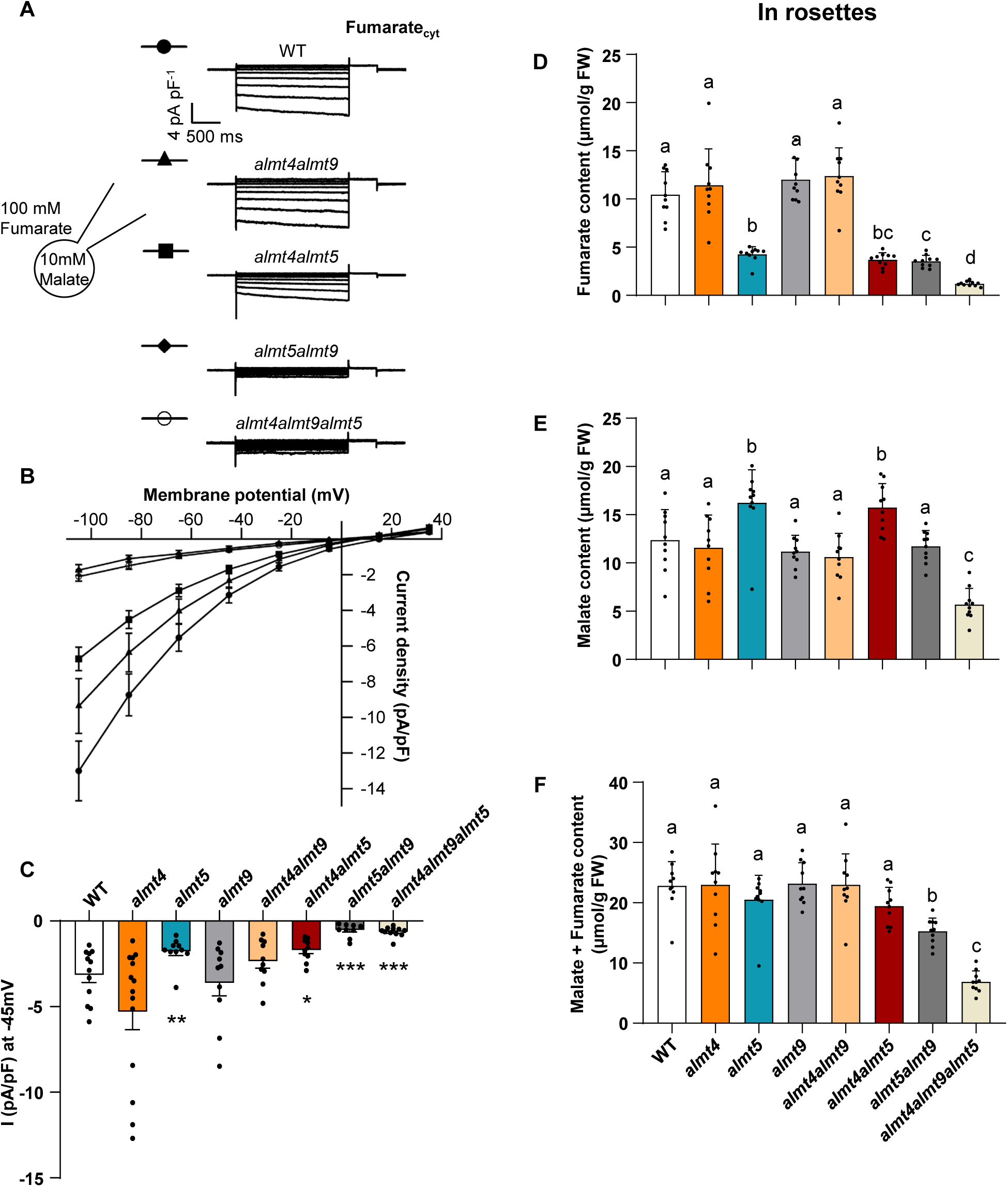
Transport capacities of the *At*ALMT channels. (**A**), Representative whole-vacuole current traces in presence of cytosolic fumarate from WT, *almt4almt9*, *almt4almt5*, *almt5almt9* and *almt4almt9almt5* plants. Starting from 0 mV, 2 second voltage steps were applied starting from +35 until - 125 mV in -20 mV decrements, each followed by a step at +35 mV for 500 ms. Holding potential was set at 0 mV. (**B**) I/V curves of the mean current densities in cytosolic fumarate condition. (**C**), Mean current densities at -45 mV from in cytosolic fumarate condition (WT: n=12; *almt4*: n=14; *almt5*: n=10; *almt9*: n=10; *almt4almt9*: n=10; *almt4almt5*: n=10; *almt5almt9*: n=10; *almt4almt9almt5*: n=11). (**B** and **C**) Data are represented as mean ± SEM. (**D-F**), Organic anion contents measured by High Pressure Ion Chromatography (HPIC) in rosettes of WT, *almt4*, *almt5*, *almt9*, *almt4almt9*, *almt4almt5*, *almt5almt9* and *almt4almt9almt5*. (**D**), Fumarate; E, Malate; F, sum of Malate + Fumarate. Plants were 40 days old grown in soil in short days conditions (8h light), and entire rosettes were collected at the end of the day (ED). Values are mean + SD, n = 10 rosettes for each genotypes. Statistical analysis : Kruskal-Wallis test + Mann-Whitney post-hoc test, Benjamini- Hochberg correction.

The fumarate^2-^ current densities measured at -45 mV in vacuoles from *almt4* (−5.3± 1.0 pA.pF^-1^) and *almt9* (−3.6 ± 0.7 pA.pF^-1^) mutants overlapped with the currents measured in wild-type (−3.15 ± 0.44 pA.pF^-1^) (Fig. 2C and Fig. S4A-B). Differently, all the mutants lacking *At*ALMT5, had drastically reduced current densities (Fig. 2A-C): *almt4almt5* (−1.69 ± 0.21 pA.pF^-1^), *almt5almt9* (−0.52 ± 0.12 pA.pF^-1^) and *almt4almt9almt5* (−0.63 ± 0.09 pA.pF^-1^). These values corresponded to a 46%, 83% and 80% decrease, respectively, compared to wild-type vacuoles (Fig. 2A-C). The malate^2-^ current densities at -47 mV were not significantly different from wild-type vacuoles in *almt4almt5* (−0.52 ± 0.06 pA.pF^-1^), *almt5almt9* (−0.43 ± 0.06 pA.pF^-1^) and *almt4almt9almt5* (−0.63 ± 0.1 pA.pF^-1^) (Fig. S4C-E). Only in *almt4almt9* (−0.33 ± 0.03 pA.pF^-1^), the malate^2-^ current density at -47 mV was slightly, yet significantly, reduced (Fig. S4E). The malate^2-^ currents in all mutants knocked out for *AtALMT9* presented instantaneous currents (Fig. S4C). Importantly, in the *almt4almt5* mutant, where only *AtALMT9* is expressed, a time-dependent activation kinetics similar to the wild-type vacuoles was observed (Fig. S4C). Overall, this patch-clamp analysis revealed that the *almt5almt9* and *almt4almt9almt5* mutants displayed dramatically reduced capacity to transport fumarate^2-^ across the tonoplast of mesophyll cells.

To understand whether the reduced fumarate transport capacities across the tonoplast impaired the accumulation of organic acids in the leaves, We quantified the content of organic acids in the rosettes sampled at the end of the photoperiod, when their concentration is maximal (Gibon *et al*., 2009; Fig. 2D-E; Supporting Table 1). Like previously observed (Doireau *et al*., 2024), compared to the wild-type (fumarate^2-^ = 10.4 ± 2.4 µmol/g FW, malate^2-^ = 12.7 ± 3.2 µmol/g FW), in *almt5* mutant, the fumarate^2-^ content was decreased by 59 % (4.3 ± 0.8 µmol/g FW) while malate^2-^ content was increased by 31% (16.2 ± 3.4 µmol/g FW) (Fig. 2D). In all the knock-out mutants lacking *AtALMT5,* fumarate^2-^ content was reduced*: almt4almt5* (3.7 ± 0.7 µmol/g FW), *almt5almt9* (3.5 ± 0.6 µmol/g FW) and *almt4almt9almt5* (1.2 ± 0.2 µmol/g FW). This corresponded to a reduction by 64%, 66%, 88% respectively in comparison to wild-type (10.4 ± 2.4 µmol/g FW). In *almt4almt5* the malate^2-^ content was 27 % increased (15.7 ± 2.5 µmol/g FW) in comparison to wild-type (12.4 ± 3.2 µmol/g FW). However, in *almt5almt9* the malate^2-^ level (11.7 ± 1.7 µmol/g FW) was equivalent to the wild-type, while in *almt4almt9almt5* it was 54% reduced (5.7 ± 1.7 µmol/g FW). The sum of malate^2-^ and fumarate^2-^ was reduced by 15% in *almt4almt5* (19.5 ± 3.1 µmol/g FW), by 33% in *almt5almt9* (15.3 ± 2.2 µmol/g FW) and by 70% in *almt4almt9almt5* (6.9 ± 1.8 µmol/g FW) in comparison to the wild-type (22.8 ± 4.0 µmol/g FW) (Fig. 2F). Citrate^3-^ level was reduced in *almt4almt9almt5* only, being 2 times lower than in wild-type (Supporting Table 1).

**Table 1:** Carbon (C) and Nitrogen (N) contents 5h after the beginning of the photoperiod.

| | %C | $\delta^{13}\text{C}$ | A%$^{13}\text{C}$ | %N |
| --- | --- | --- | --- | --- |
| WT | 33,44 ± 0,92 | -31,35 ± 0,32 | 1,12 ± 0,00 | 6,35 ± 0,14 |
| *almt5* | 33,49 ± 1,18 | -31,41 ± 0,31 | 1,12 ± 0,00 | 6,26 ± 0,32 |
| *almt9* | 33,97 ± 0,89 | -31,41 ± 0,18 | 1,12 ± 0,00 | 6,26 ± 0,17 |
| *almt4* | 33,95 ± 0,99 | -31,32 ± 0,29 | 1,12 ± 0,00 | 6,38 ± 0,18 |
| *almt5almt9* | **34,76 ± 1,09 \*\*** | **-29,86 ± 0,33 \*\*\*** | 1,12 ± 0,00 | **6,15 ± 0,19 \*** |
| *almt4almt5* | 34,18 ± 0,94 | -31,58 ± 0,26 | 1,12 ± 0,00 | **6,08 ± 0,12 \*\*** |
| *almt4almt9* | 34,31 ± 1,02 | -31,44 ± 0,18 | 1,12 ± 0,00 | **6,13 ± 0,19 \*** |
| *almt4almt9almt5* | **35,16 ± 0,93 \*\*\*** | **-27,86 ± 0,38 \*\*\*** | 1,12 ± 0,00 | **5,89 ± 0,20 \*\*\*** |

Carbon and Nitrogen content measurement of WT, *almt4*, *almt5*, *almt9*, *almt4almt9*, *almt4almt5*, *almt5almt9* and *almt4almt9almt5* 40 days-old rosettes grown in short days photoperiod (8h light). Entire rosettes were collected 5h after the beginning of the photoperiod in the growth chamber. Each measurement correspond to n = 10 rosettes. Data were obtained by mass spectrometry. Data are mean ± SD. Asterisks show a difference from the WT. Statistical analysis : Mann-Whitney test (\*P<0.05;\*\*P<0.01;\*\*\*P<0.001).

Interestingly, the levels of the major inorganic anions (chloride, sulphate, phosphate and nitrate) were not significantly modified in vacuolar ALMT mutants (Supporting Table 1). Overall, the reduction of organic acid levels in the different mutants correlates with the reduction of organic acid transport capacities observed by patch-clamp analysis and demonstrates that *At*ALMT5, *At*ALMT4 and *At*ALMT9 are key players for the transport of fumarate^2-^ and malate^2-^ in the vacuoles of mesophyll cells. However, the role of the three ion channels is not equivalent. Indeed, *At*ALMT4 appears to have a minor role while *At*ALMT5 displays a specific function in fumarate^2-^ transport. Finally, *At*ALMT9 is instead involved in both malate^2-^ and fumarate^2-^ transport.

### Altered K^+^ levels and cell sap pH in *AtALMT* mutants

The levels of anionic and cationic species in plant cells are essential to keep the electroneutrality of the cellular solution and to maintain the turgor pressure. In order to evaluate to which extent a reduction of organic anion content can be compensated by a reduction of cation levels, the major inorganic cations (K^+^, Mg^2+^, Ca^2+^, Mn^2+^, Zn^2+^, Fe, Cu) and total sulphur and phosphorus content and were quantified in whole rosettes of all mutant lines (Fig. 3, S5). Despite the leaf senescence observed in *almt5almt9* and *almt4almt9almt5* mutants, no iron deficiency was detected (Fig. S5). Notably, the levels of K^+^ were strongly reduced in *almt4almt9* (44,70 ± 5,68 mg Elt/g DW), *almt4almt5* (45,72 ± 5,14 mg Elt/g DW), *almt5almt9* (47,48 ± 3,71 mg Elt/g DW) and *almt4almt9almt5* (21,29 ± 4,58 mg Elt/g DW) corresponding to a decrease by 32%, 31%, 28% and 68% compared to wild-type (66,0 ± 6,3 mg Elt/g DW), respectively. In the single mutants we found only a moderate K^+^ reduction of 14.6% in *almt4* (56,4 ± 4,8 mg Elt/g DW) and of 15.8% in *almt9* (55,6 ± 6,3 mg Elt/g DW). Then the expression level of the two major vacuolar K^+^ transporters, *AtNHX1* and *AtNHX2,* was quantified in the multiple mutants (Bassil *et al*., 2011; Barragán *et al*., 2012; Fig. 3). Only the expression of *AtNHX2* was modified in the *almt4almt9almt5* being 44% lower than in the wild-type (Fig. 3C) indicating that the observed K^+^ decrease did not depend on changes of the expression of *AtNHX1* and *AtNHX2*. The transport of potassium and the subcellular concentrations of malate^2-^ and fumarate^2-^ are part of the machinery controlling cellular pH homeostasis (Arias *et al*., 2013). Therefore, the cell sap pH from leaves collected at the end of the day was quantified in the single and multiple *AtALMT* mutants (Fig. 3D). A significant increase of the cell sap pH was observed in both *almt5almt9* (6.07 ± 0.05) and *almt4almt9almt5* (6.24 ± 0.06) in comparison to wild-type (5.99 ± 0.08). Conversely, *almt4*, *almt5*, *almt9* and the double mutants *almt4almt9* and *almt4almt5* did not show cell sap pH difference. Thus, the reduction of K^+^ was not sufficient to compensate the extreme reduction of organic anions in the triple mutant. Overall, the reduced capacities to store malate and fumarate in the vacuole have an important influence on the whole cell physiology by modifying key biochemical parameters.

**Figure 3.**
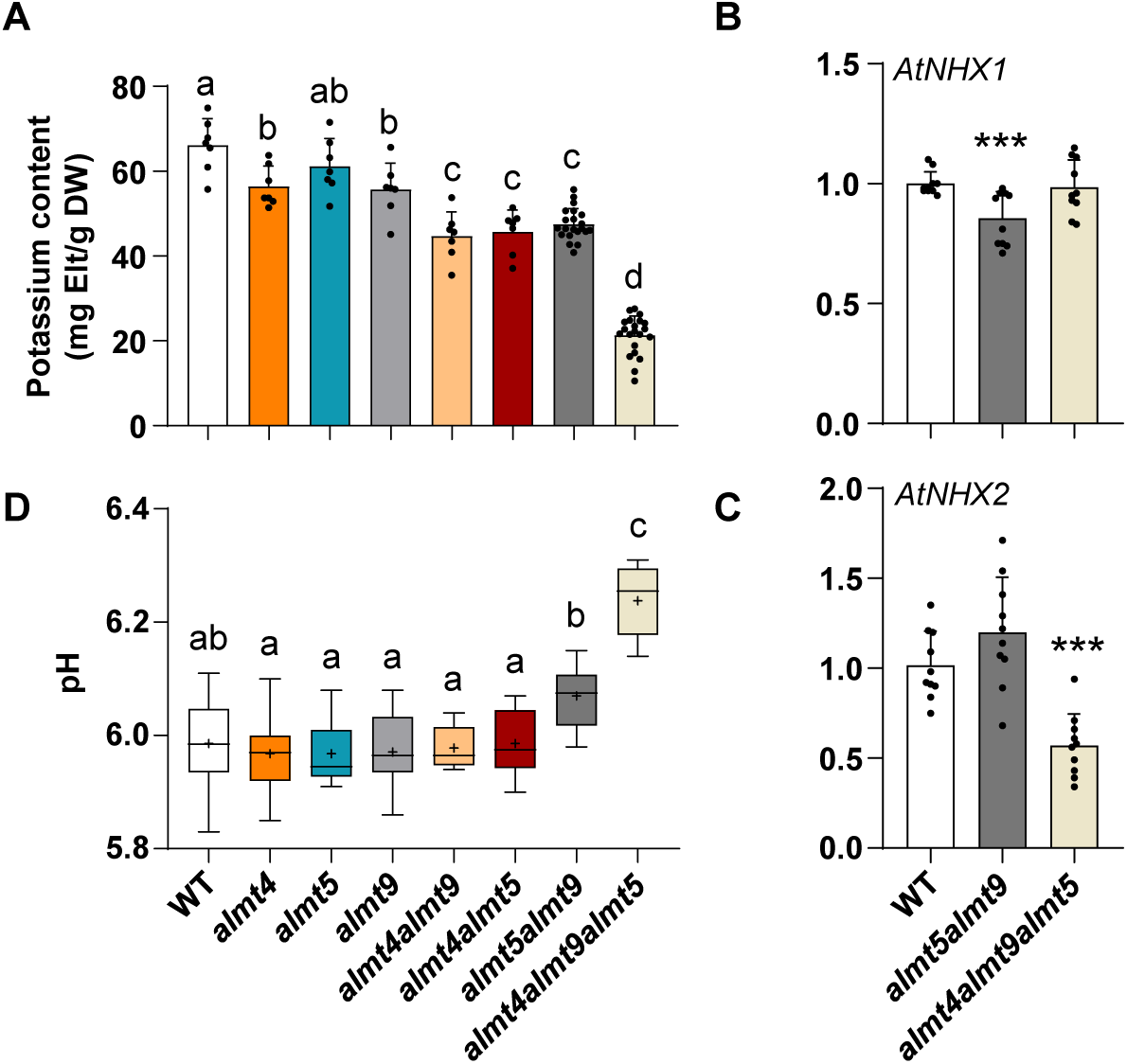
: pH and potassium/ potassium gene expression modifications. (**A**) Potassium content of entire rosettes measured by inductively coupled plasma optical emission spectrometer (ICP-OES). Plants were grown in short days conditions (8h light) and entire rosettes were collected at the end of the day (ED). Value are mean + SD of n= 7-21 rosettes for each genotypes. Asterisks show the difference from the WT. Statistical analysis : Kruskal-Wallis test + Mann-Whitney post-hoc test, Benjamini-Hochberg correction. (**B,C**) *AtNHX1* (**B**) and *AtNHX2* (**C**) expression quantified by qRT-PCR. 40 days-old plants were grown in short days conditions (8h light). Leaves number 9 were collected at the end of the day (ED). For each genotype 10 plants were collected and analysed pooled by two. Transcript accumulation was quantified after normalisation by the reference gene Yellow leaf specific gene (YLS8). Statistical analysis: Mann-Whitney test (*P<0.05; **P<0.1; ***P<0.001). (**D**) The loss of *almt5almt9* and *almt4almt9almt5* increases the cell sap pH in the plant rosettes. Plants were grown 40 days in short days conditions (8h light). Box plots with min/max whiskers represents the distribution of 10 plants per genotypes. Asterisks show the difference from the WT. Statistical analysis : Kruskal-Wallis test + Mann-Whitney post-hoc test, Benjamini-Hochberg correction.

### An impaired shoot organic acid accumulation modifies carbon and nitrogen metabolism

Since malate^2-^ and fumarate^2-^ are key intermediates of plant central carbon metabolism and precursors of carbon skeletons for nitrogen assimilation, the functioning of the central carbon and nitrogen metabolisms has been studied in the mutant lines. First, the absolute quantification of carbon and nitrogen pools was performed in all mutant lines using EA-IRMS (Table 1 and Supporting Table 2). The data showed a significant increase of the total C content (%C) in both *almt5almt9* (+1.3 %) increase, and *almt4almt9almt5* (+1.7 %) increase, compared with the wild-type (Table 1). In both *almt5almt9* and *almt4almt9almt5*, the natural ^13^C abundance (δ^13^C) was significantly higher than in the wild-type. Given the reduced FW/DW ratio in these mutants (Fig. 1D), this isotopic discrimination could reflect both a higher water use efficiency (Pater *et al*., 2017; Yu *et al*., 2024) and a modification of metabolic fluxes operating isotopic fractionation (Tcherkez *et al*., 2011). In the complemented lines *almt5almt9/gALMT5* and *almt5almt9/gALMT9*, the total C content and the δ^13^C were like in the wild-type, confirming the link between vacuolar *At*ALMT channels and the modification of C contents (Supporting Table 2). In the *almt5almt9*, *almt4almt5*, *almt4almt9* and *almt4almt9almt5*, the total nitrogen (%N) in the rosettes was significantly reduced compared to the wild-type (Table 1). This could reflect a reduction of N assimilation due to the limitation of organic acid availability. Taken together, these data show a global modification of plant metabolism in the two mutants, *almt5almt9* and *almt4almt9almt5* showing a reduced organic acid storage (Fig. 2F), and biomass (Fig. 1).

Since carbon and nitrogen metabolism appear to be modified in *almt5almt9* and *almt4almt9almt5*, an evaluation of the functioning of plant central metabolism was achieved using transient ^13^CO2-labelling of mutant lines. For this purpose, the rosette of wild-type, *almt5*, *almt9*, *almt4*, *almt5almt9* and *almt4almt9almt5* mutant lines was integrated into an illuminated custom chamber to deliver 99% ^13^CO2 for 4h during the photosynthetic period (Fig. S1). ^13^C-enrichment and absolute content of sugars, organic and amino acids were quantified by GC-MS and converted to ^13^C allocation fluxes (Fig. 4). Overall, all the genotypes incorporated similar amounts of ^13^C during the experiment (Fig. S6), thus confirming that differences of ^13^C allocation into metabolites will reflect differences of metabolic fluxes. In wild type plants, the free soluble metabolites analysed by GC-MS accounted for nearly 13% of the net ^13^C fixed (Fig. 4A), consistent with the fact that the majority of C is fixed into starch in the light. Interestingly, in the wild-type Arabidopsis, a non-negligible part of the photosynthetic carbon is incorporated into fumarate^2-^ (4%) and malate^2-^ (2%) compared to sugars (2.5%) and amino acids (4%) (Fig. 4A). Thus, fumarate^2-^ and malate^2-^ likely represent important secondary stocks of carbon in mesophyll cells. In *almt5almt9* and *almt4almt9almt5* we observed a drastic reduction of ^13^C incorporated into fumarate^2-^, being 2% and 0.5%, respectively. This result confirmed that *de novo* biosynthesis of fumarate^2-^ was partially controlled by the capacity of mesophyll cells to store fumarate within vacuoles. Interestingly, the ^13^C that is not incorporated into fumarate^2-^ and malate^2-^ is not redirected to major pools of soluble sugars like glucose, fructose and sucrose (Fig. 4C). The results showed that in *almt5almt9* and *almt4almt9almt5*, assimilated ^13^CO2 may be preferentially redirected to the forward branch of the TCA cycle and *de novo* nitrogen assimilation (Fig. 4B). Indeed, the ^13^C allocation to citrate^3-^, α-ketoglutarate (2-OG) and glutamate was significantly increased in the *almt5almt9* and *almt4almt9almt5* genotypes compared to the wild-type (Fig. 4C). Since the carbon skeleton of glutamate derives from the citrate-2-OG branch of the TCA cycle (Sweetlove *et al*., 2010), our results suggested a higher flux for this branch in the double and triple mutants. However, the cyclic mode of the TCA cycle was still maintained in these mutants, as evidenced by the maintenance of the ^13^C allocation into succinate compared to the wild-type (Fig. S7). Hence, in these mutants, part of the carbon that is not escaping the TCA cycle through fumarate is likely to escape as 2-OG and could support nitrogen assimilation. In line with this, ^13^C allocation to glycine, a photorespiratory intermediate, was increased both in *almt5almt9* and *almt4almt9almt5* mutants, but not for serine (Fig. 4C). This could reflect an inhibition of the mitochondrial glycine decarboxylase activity through NADH production by TCA cycle dehydrogenases (Bykova *et al*., 2005). Finally, the analysis of carbon isotopologue distribution revealed a higher production of fully labelled malate^2-^, fumarate^2-^ and citrate^3-^ in the double and triple mutants compared to the wild-type, thereby supporting a higher turn-over for the TCA cycle. However, fumarate displayed a different pattern of carbon isotopologue distribution compared to succinate in all Arabidopsis lines (Fig. S8). Thus, malate^2-^ and fumarate^2-^ accumulation in all lines still relies significantly on the cytosolic PEPc activity, independently of the mitochondrial TCA cycle.

**Figure 4:**
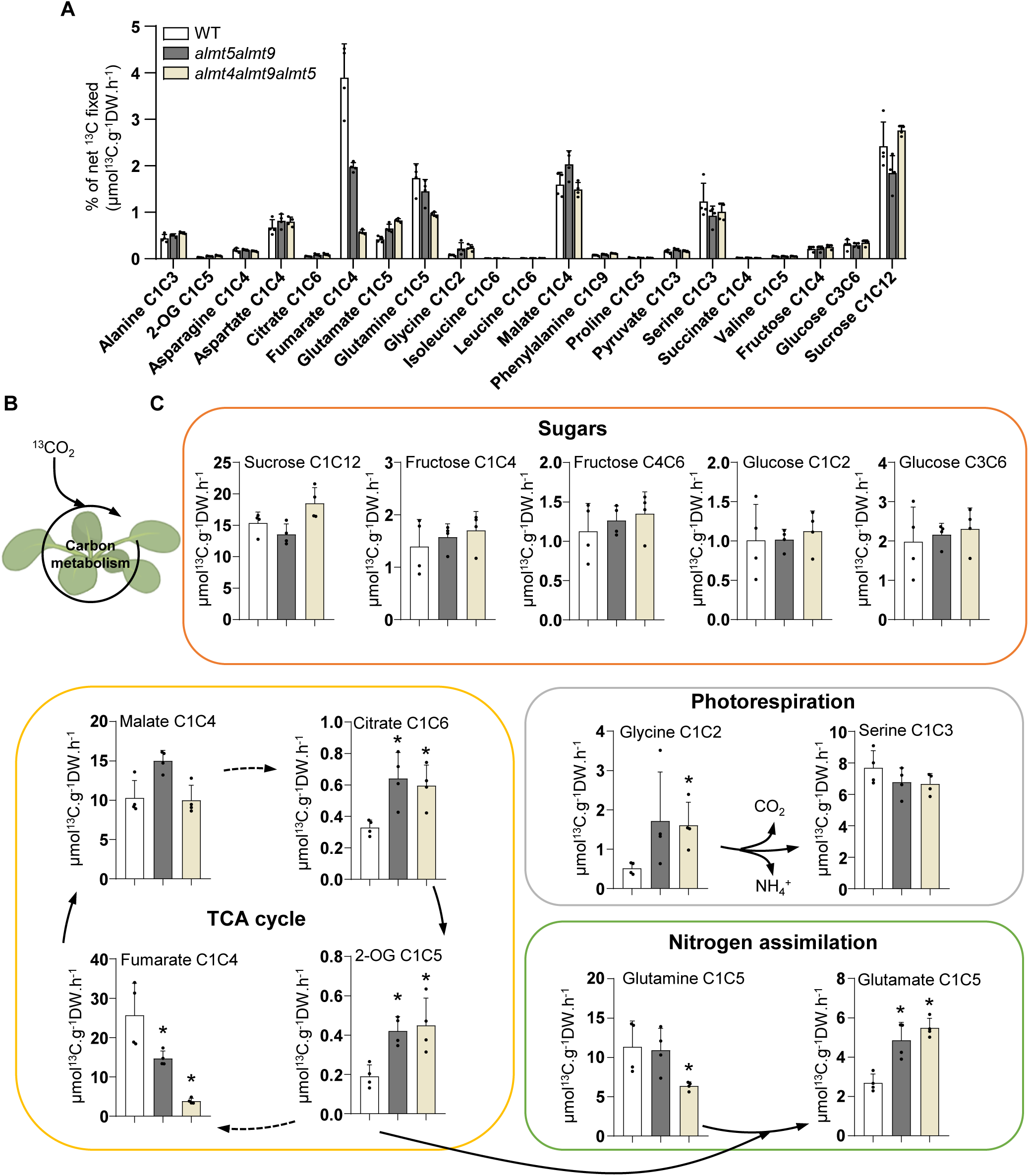
^13^CO_2_ incorporation in *Arabidopsis thaliana* rosettes metabolism. (**A**), Metabolites percentage of net ^13^CO_2_ enrichment of WT, *almt5almt9* and *almt4almt9almt5* plant after 4h of ^13^CO_2_ incorporation (n=4 entire rosettes). Plants were 6-week-old grown in short days (8h light) and placed in a ^13^CO_2_ enriched chamber 2-3h after the beginning of the light period. (**B**), ^13^CO_2_ is incorporated in the plant carbon metabolism and serves as carbon skeletons during the synthesis of different metabolites. (**C**), ^13^C incorporation quantification in metabolites and amino acids on WT, *almt5almt9* and *almt4almt9almt5* (n = 4 entire rosettes). Quantification was done by GC-MS after 4h of ^13^CO_2_ incorporation. Statistical analysis: Mann-Whitney test (*P<0.05). Asterisks represents a significant difference compared to the WT. Results are expressed in µmol^13^C.g^-1^DW.h^-1^.

**Figure 9.**
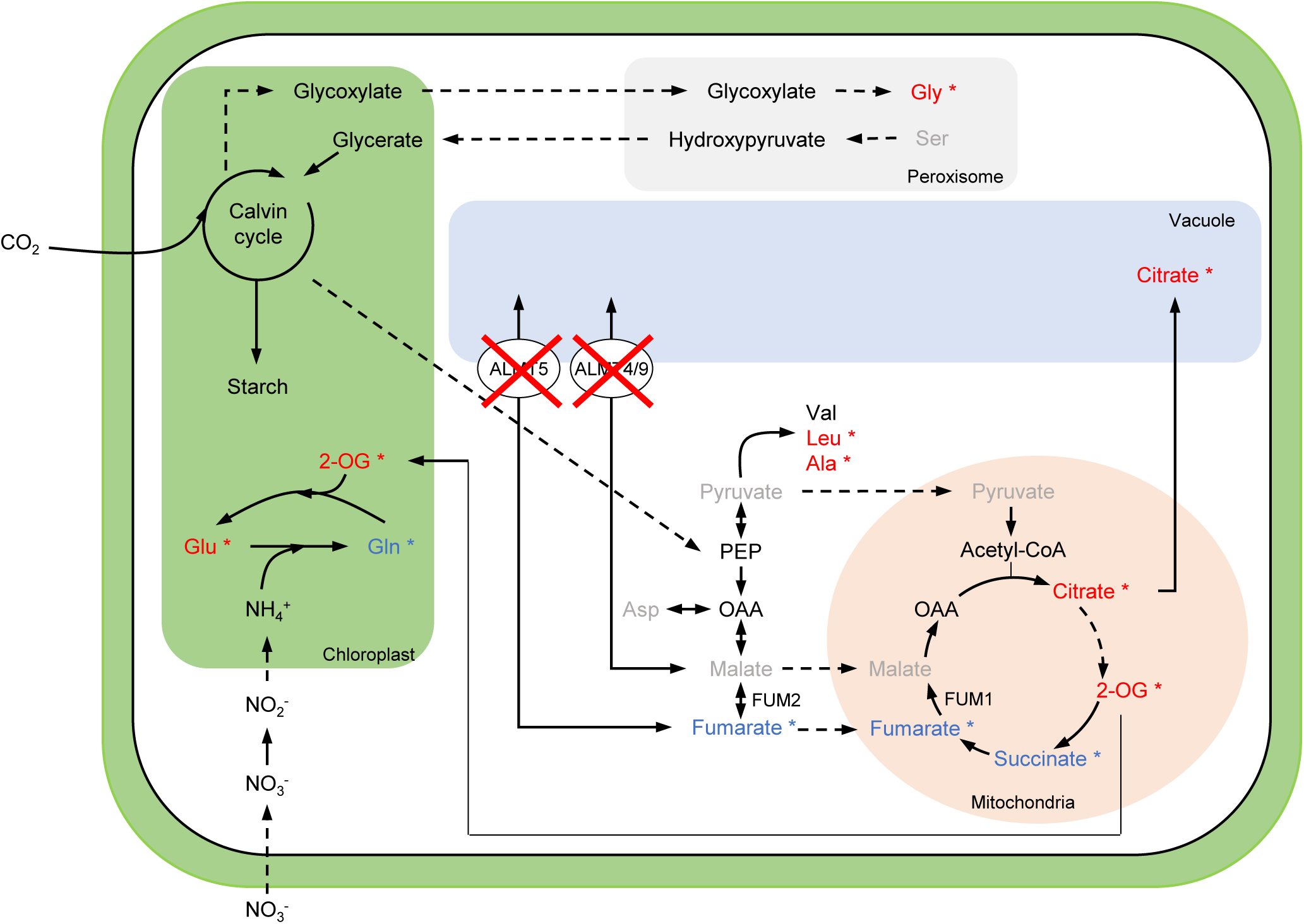
Metabolites changes in *almt4almt9almt5*. Metabolites data from the ^13^C experiment described in Fig. 9 in a metabolism map. Solid arrows represent single enzymatic reaction, and dashed arrows represent multiple reactions or transport of metabolites from one compartment to another. Colours indicate significant metabolites increase (*red*) or decreased (*blue*), or unchanged (*gray*). Metabolites than weren’t tested are written in black.

## DISCUSSION

Land plants store a significant part of the carbon fixed by photosynthesis under the form of organic acids (i.e; malate^2-^ and fumarate^2-^). These ions are generally stored in the vacuoles and remobilised depending on the photoperiod (Ludwig *et al*., 2024). In C3 plants, representing the vast majority of terrestrial plants, both malate^2-^ and fumarate^2-^ are stored into the vacuole during daytime but the metabolic importance of this process remains to be addressed. For instance, most of the mutant lines characterised in Arabidopsis for vacuolar transporters of malate^2-^ and fumarate^2-^ do not show a growth phenotype (Hurth *et al*., 2005; Meyer *et al*., 2011; De Angeli *et al*., 2013b; Eisenach *et al*., 2017; Frei *et al*., 2018; Ye *et al*., 2021; Doireau *et al*., 2024). However, the vacuolar transporters mediating malate^2-^ and fumarate^2-^ transport were studied separately. Notably, there are three ALMTs transporters expressed in the mesophyll cells, encoding proteins addressed to the tonoplast and with possible redundant functions (Kovermann *et al*., 2007; Meyer *et al*., 2011; Eisenach *et al*., 2017; Doireau *et al*., 2024). Combining mutations of different genes within the ALMTs family can be an interesting strategy to massively disrupt vacuolar transport of organic acids and further investigate the associated metabolic consequences.

In this study, we disrupted the transport of organic acids to the vacuole by generating multiple knock-out of the vacuolar *At*ALMTs (*At*ALMT4, *At*ALMT5 and *At*ALMT9) expressed in the mesophyll cells, the major photosynthetic tissue. The electrophysiological analysis of the vacuolar transport capacity showed that in *almt5almt9* and *almt4almt9almt5* mutants, fumarate^2-^ and malate^2-^ ionic currents were dramatically impaired (Fig. 2; Fig. S4). In line with this result, these two mutants lost their capacity to accumulate fumarate^2-^ in the leaves, and in *almt4almt9almt5,* malate^2-^ loading was reduced too (Fig. 2). Therefore, these mutants provide the opportunity to evaluate the impact of abolishing fumarate^2-^ and malate^2-^ transport to the vacuole in a C3 plant. Notably, *almt5almt9* and *almt4almt9almt5,* the mutants with abolished malate^2-^ and fumarate^2-^ transport to the vacuole, displayed a dramatic growth defect (Fig. 1A). Collectively, these data demonstrate that in Arabidopsis, a C3 plant, the vacuolar storage of these organic acids is necessary for growth and biomass production.

Malate^2-^, and fumarate^2-^ in Arabidopsis, are organic acids sitting at the crossroad of several metabolic pathways involved in carbon fixation and amino acid synthesis processes (Mathieu *et al*., 1986; Hurth *et al*., 2005; Fernie & Martinoia, 2009). Further, these two organic acids are also involved in the regulation of ionic, redox and pH homeostasis in plant cells (Meyer *et al*., 2010). The dramatic growth phenotype we observed could result from the alteration of these processes. Interestingly, *almt5almt9* and *almt4almt9almt5*, the two mutants presented major growth defects and displayed a higher cell sap pH (Fig. 3D). In *almt4almt9almt5* the cell sap pH is comparable to the one observed in the vacuolar proton mutants *vha2vha3* (Krebs *et al*., 2010), suggesting that malate^2-^ and fumarate^2-^ accumulation significantly participate to the vacuolar pH homeostasis. Similarly, in CAM plants malate^2-^ accumulation and proton pumps are functionally linked (Winter & Smith, 2022; Hafke et al. 2003). Further, *almt5almt9* and *almt4almt9almt5* also display a lower K^+^ content in the shoots (Fig. 3A). The Arabidopsis mutant for the vacuolar K^+^ transporter NHX1 also exhibited reduced K^+^ content and delayed growth compared to the wild-type or complemented lines (Apse *et al*., 2003). However, we showed here that the K^+^ under accumulation was not due to reduced expression of NHX1 and NHX2, the two main vacuolar K^+^ transporters (Fig. 3B-C). In addition, in fruits, potassium nutrition has a direct effect on malate^2-^ accumulation (Zhang *et al*., 2018). Our data show that within the leaves the reduced capacity to transport malate^2-^ and/or fumarate^2-^ in the vacuole influence has an impact on potassium accumulation. These results are demonstrating a direct link between malate/fumarate accumulation and potassium content (Fig. 2D-F; Fig. 3A). Given the important contribution of potassium to the osmotic potential of plant cells (Poucet *et al*., 2022; Boulc’h *et al*., 2024), a reduced content of this ion and of fumarate^2-^ or malate^2-^ likely directly influence circadian variations of turgor, and consequently cell expansion and plant growth (Ali *et al*., 2023). Besides this, knock-out mutants for two Cl^-^ anion channels in Arabidopsis also showed a reduced vacuolar ion content combined with delayed root-hair elongation (Zhang *et al*., 2017). Thus, this mechanism may not be restricted to the imbalance of one particular ion within the vacuole and should partially rely on the contribution of organic ions.

Interestingly, up to 4% of the ^13^CO2 was incorporated into fumarate^2-^ in the wild-type plants (Fig. 4A), hence confirming that fumarate^2-^ is an important secondary pool of carbon stocks in Arabidopsis cells as for soluble sugars. A ^13^CO2-labelling experiment on Arabidopsis rosette during 5 hours also found that fumarate and soluble sugars represented the major part of ^13^C- labelled soluble metabolites (Dellero *et al*., 2016). Therefore, disrupting the vacuolar storage capacity of plants was expected to have important metabolic consequences for C assimilation and its use. Yet all mutant lines showed similar net ^13^C fixed, but the variation of δ ^13^C values suggested contrasted metabolic functioning in *almt5almt9* and *almt4almt9almt5* (Fig. S6, Table 1). ^13^C allocation patterns and isotopologue distribution within TCA cycle intermediates showed that fumarate was preferentially synthesised from PEPc over the TCA cycle in *almt5almt9* and *almt4almt9almt5.* Interestingly, the overexpression of PEPc in a C3 plant significantly increased δ ^13^C values of whole dry matter (Giuliani *et al*., 2019). This higher flux for PEPc would also stimulates in return the *de novo* production of citrate from TCA cycle, especially in a context of disrupted vacuolar storage of both malate and fumarate. In the *almt5almt9* and *almt4almt9almt5* mutants, ^13^CO2 incorporation into citrate^3-^ and 2-OG was higher, (Fig. 4C) and confirmed a preponderant non-cyclic TCA behaviour during daytime (Sweetlove *et al*., 2010). Further evidence for the flux modes are highlighted by the reduced conversion of glycine into serine by the glycine decarboxylase (GDC) based on ^13^C allocation patterns. This inhibition has been attributed to the concurrent use of NAD by TCA cycle dehydrogenase and GDC in the mitochondria and/or excess NADH production (Bykova *et al*., 2005). This flux mode is known to support *de novo* nitrogen assimilation through GS-GOGAT cycle and to actively contribute to the release of carbon from the TCA cycle (Dellero *et al*., 2023). In line with this, in both *almt5almt9* and *almt4almt9almt5* the accumulation of ^13^C into glutamate was increased (Fig. 4C). However, ^13^C allocation to glutamine was decreased in the meantime and correlated with the lower amount of total N observed in these mutants compared to the wild-type (Fig. 4, Table 1). In addition, low N content in *almt5almt9* and *almt4almt9almt5* mutants was associated with the presence of senescing leaves with a strong expression of *SAG13* (Table 1, Fig. S1). These phenotypes suggest an important role of the vacuole in the regulation of cytosolic malate^2-^ (and fumarate^2-^) levels for the functioning of plant central metabolism. Indeed, the malate valve is a key regulator of the redox potential of subcellular compartments and partially fuels nitrate reduction with NADH (Selinski & Scheibe, 2019). In tobacco, the manipulation of NAD/NADH levels has a great impact on N assimilation (Dutilleul *et al*., 2005). Therefore, N assimilation in these mutants might be partially constrained by redox-dependent reduction of nitrate to ammonia rather than the production of C skeletons by the TCA cycle. Besides this, the vacuole was recently identified as an important player of photorespiration through the sequestration of glycerate in response to N starvation (Lin & Tsay, 2023). This further confirm the importance of vacuolar storage of organic acids for the functioning of plant central metabolism.

In conclusion, our results demonstrate in a C3 plant that the capacity to transport organic acids like malate^2-^ and fumarate^2-^ in the vacuole of mesophyll cells is essential for plant growth. Indeed, our results show that malate^2-^ and fumarate^2-^ transport across the vacuolar membrane is an essential step to regulate metabolic processes but also pH and ion homeostasis within plant cells. In CAM plants, malate^2-^ transport in the vacuole is essential for this specialised metabolism adapted to warm and dry environment. CAM metabolism evolved independently several times for C3 plants (Edwards, 2019). Our data demonstrate that also in C3 plants vacuolar transport of malate^2-^ and fumarate^2-^ is essential for the regulation of plant metabolism and growth. It is tempting to speculate that evolution of CAM exploited this pre- existing property coordinating it with a new metabolic organisation. So far, the vacuole transport of these organic acids in photosynthetic tissues and its importance in C3 plants was not clear. Our results identified the key molecular players of the process and indicate that this process could be a leverage to improve plant biomass production in the future.

## Acknowledgements

We thank K. Bertaux for the technical support. We thank the Plant Electrophysiology Platform (PEP) and C. Corratgé-Faille from the Institute for Plant Sciences of Montpellier (IPSiM) for technical support and assistance. We acknowledge the Isotope Quantification Platform (AQUI) and T. Perez assistance. We acknowledge the SAME platform and S. Chay for the metal and cations measurements. We thank the Metabolic Profiling and Metabolomic Platform (P2M2, https://p2m2.hub.inrae.fr/) for providing instrument devices to quantify the ^13^C enrichment of plant central metabolites. We are grateful to Dr. Anne Marmagne for performing C quantification during ^13^CO2 incorporation experiments. This work was funded by CNRS grant (ATIP-Avenir) and the French National agency of Research (grant Netflux) to ADA. RD was funded by a PhD grant from University of Montpellier and from INRAE.

## Contributions

Plant phenotyping and pictures were done by RD, EDC and ADA. Expression analyses were done by RD, data analysis was done by RD. Patch-clamp analysis were done by EDC, data analysis was done by EDC. Anion content measurements were done by RD and ADA. Plant crossing was done by PCF. Selection of knock-out mutants was done by PCF and RD. ^13^CO2 labelling experiments and GC-MS analysis were done by YD and SB, data were analysed by YD with a contribution of RD and ADA. RD and ADA wrote the manuscript with the contribution of EDC and YD. ADA and RD conceived the research.

## Data availability

All data generated and/or analyzed during the current study are available from the corresponding author on reasonable request.

