## Supporting figures and files for "Cracking vacuolar fumarate and malate transport shows its function in Arabidopsis metabolism and growth"

**Supplementary material and methods**

***In vitro* cultures**

*In vitro A. thaliana* cultures were grown in ½ MS medium supplemented with 0.05% MES buffer, 0.7% (w/v) agar, and depending on the conditions with 1% sucrose. For this, seeds were surface –sterilised and grown in a growth chamber with a day/night regime of 16h/8h, 21°C/21°C, LED lights 150 µmol.$m^{-2}$.$s^{-1}$. Seeds were stratified 72h at 4°C in the dark.

**RT-PCR**

The integrity of the *AtALMT5 and AtALMT9* mRNA was verified in the T-DNA lines (*almt5-1* and *almt5-2*) and in the multiple knock-out lines and in complemented T3 lines through RT-PCR. Leaf RNA extraction was performed by using the Direct-zol RNA Miniprep Plus kit (Zymo Research) following the manufacturer's protocol. Contaminating DNA was eliminated from the samples by RNAse-free DNAse I kit (ThermoFisher Scientific, USA), and RNases were also inhibited by using RiboLock RNase Inhibitor (Thermo Fisher Scientific). One microgram of total RNA was used as template for the first strand cDNA synthesis using the M-MLV Reverse Transcriptase enzyme (Promega) following the manufacturer's instructions. The RT-PCR was carried out using suitable primers to amplify the whole gene *AtALMT5* (*At*1g68600) (Supporting Table 4). Primers were designed using Primer3 (Rozen & Skaletsky, 2000) and OligoCalc software (Kibbe, 2007). The RT-PCR amplification was performed using the GoTaq® DNA Polymerase (Promega) and the amplification of cDNAs was arrested at the exponential phase of the PCR reaction. The Actin2 (*At*3g18780) expression level was used as loading control.

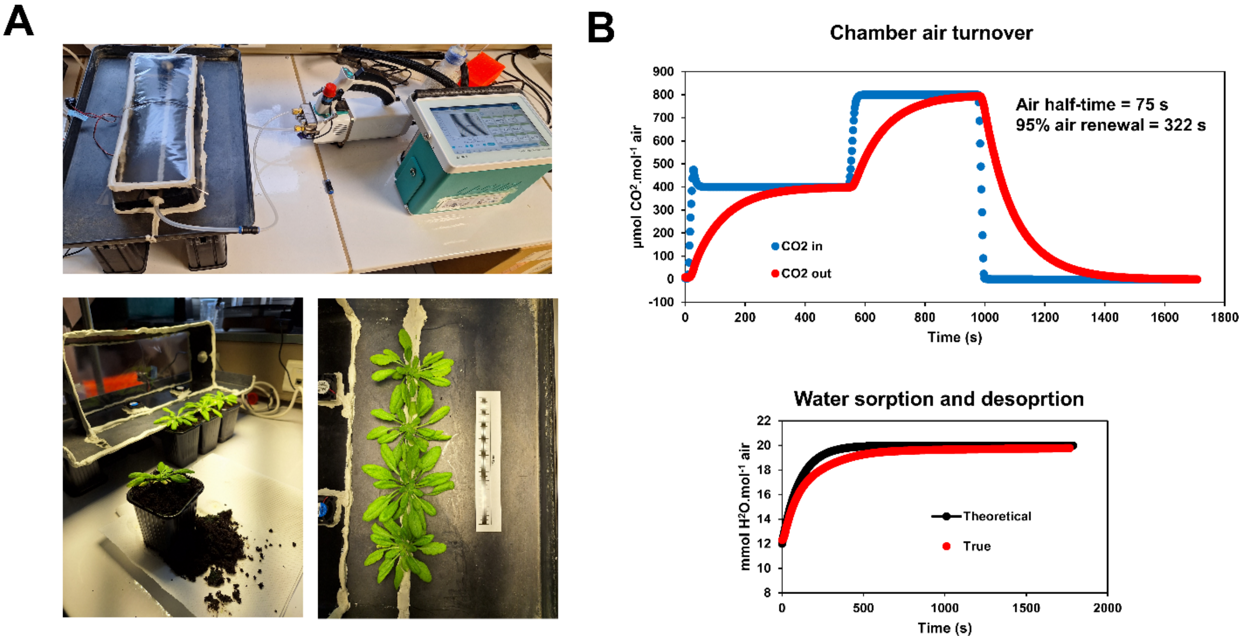

**Supporting Figure 1 : Experimental setup used to perform $^{13}CO_2$ labelling experiments.**

(A) Custom chamber connected to the LI-6800-F and integration of intact Arabidopsis rosettes.

(B) Characteristics of the custom chamber

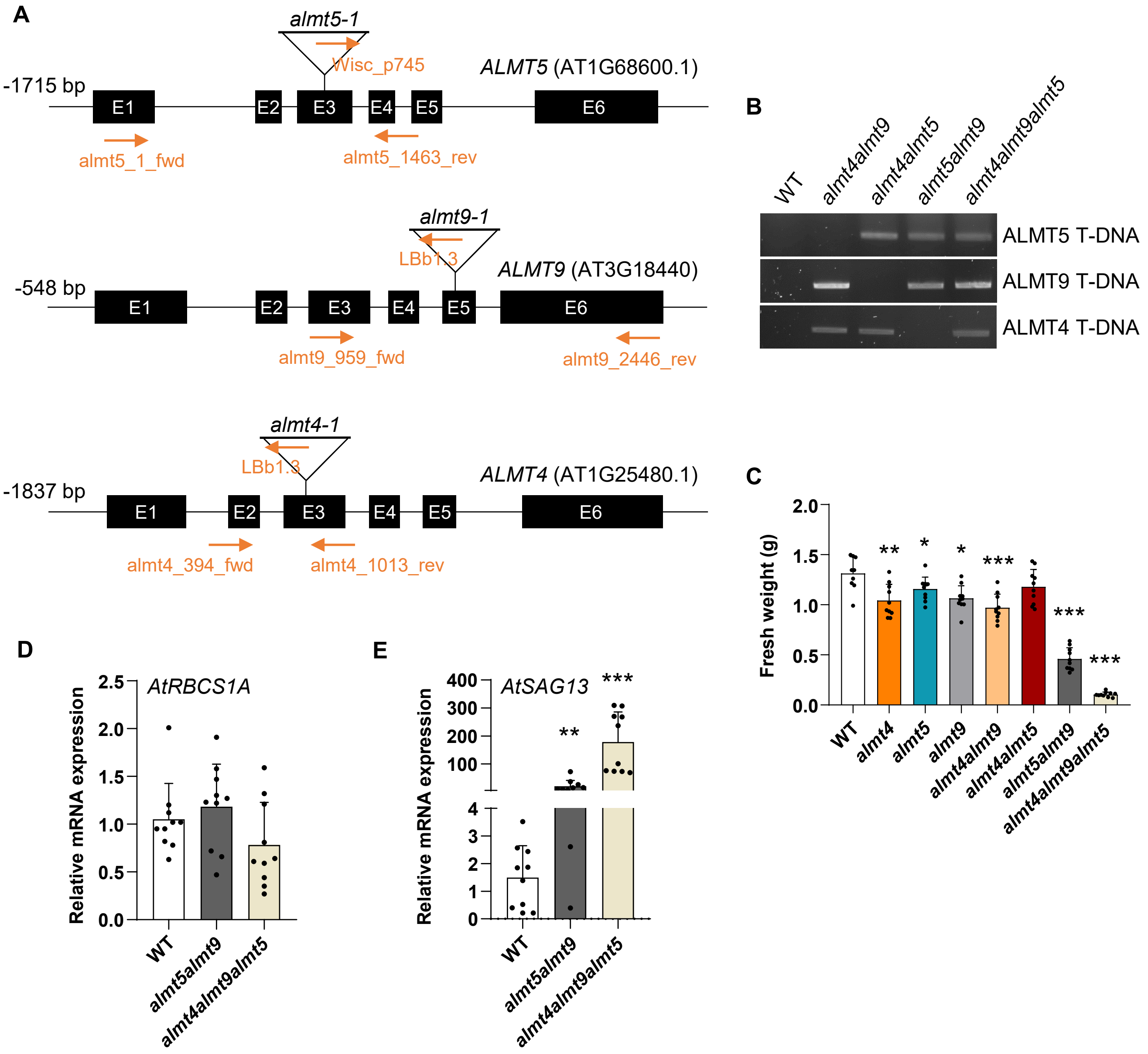

**Supporting Figure 2 : Genotype confirmation**

A, Gene structure of *AtALMT5* (At1g68600) and localisation of the T-DNA insertions in the almt5-1 (WiscDsLox386E04) transgenic lines (top). Gene structure of *AtALMT9* (At3g18440) and localisation of the T-DNA insertion in almt9-1 (Salk_055490) transgenic line (middle). Gene structure of *AtALMT4* (At1g25480.1) and localisation of the T-DNA insertion in almt4-1 (Salk_086236) transgenic line (bottom). Closed boxes represent exons (E) and solid lines represent introns. PCR amplification primers used for genotyping are indicated with arrows.

B, Verification of ALMT9, ALMT5 and ALMT4 T-DNA insertion by PCR in the positive control (WT), the doubles mutants *almt4almt9, almt4almt5, almt5almt9* and the triple mutant *almt4almt9alm5*. The primers used are the ones on T-DNA insertion in A.

C, Rosette fresh weight (FW) of *Arabidopsis thaliana* 40-days old plants knock-out lines grown in short days conditions (8h light). Values are mean + SD, n= 10 plants for each genotype. Asterisks indicate a significant difference in comparison to WT. Statistical analysis : Mann-Whitney test, (*P<0.05;**P<0.01;***P<0.001).

(D,E) *AtRBCS1A* (D) and *AtSAG13* (E) expression quantified by qRT-PCR. 40 days-old plants were grown in short days conditions (8h light). Leaves number 9 were collected at the end of the day (ED). For each genotype 10 plants were collected and analysed pooled by two. Transcript accumulation was quantified after normalisation by the reference gene Yellow leaf specific gene (YLS8). Statistical analysis: Mann-Whitney test (**P<0.1; ***P<0.001).

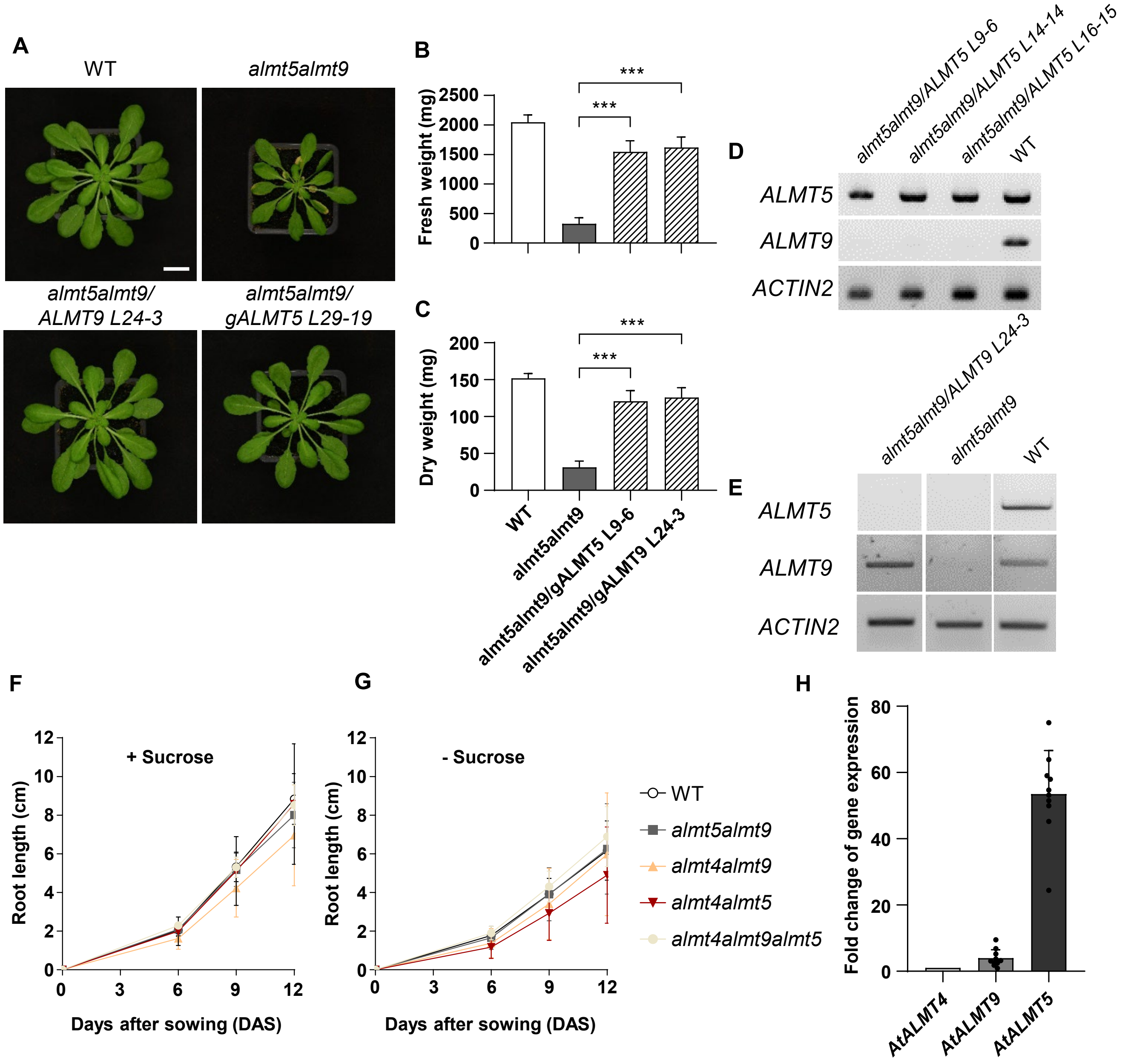

**Supporting Figure 3 : Complementation of *almt5almt9* and root growth of the different mutants**

A, *Arabidopsis thaliana* wild-type (WT), *almt5almt9* and complemented lines with ALMT5 (*almt5almt9/gALMT5*) and ALMT9 (*almt5almt9/gALMT9*) genomic fragment. Plants were 40 days old grown in short days (8h light). Bar = 2cm.

(B,C) Fresh weight (B) and dry weight (C) of WT, *almt5almt9* and complemented lines with ALMT5 (*almt5almt9/gALMT5*) and ALMT9 (*almt5almt9/gALMT9*) genomic fragment. Plants were 40 days old grown in short days (8h light). Statistical test : Mann Whitney (***P<0.001).

(D-E), *At*ALMT5 (D) and *At*ALMT9 (E) transcript determined by Reverse Transcriptase (RT)-PCR analysis of total RNA isolated from leaves of complemented plants in comparison to the control (WT). The *AtACTIN2* transcript was used as control of RNA loading.

(F,G) Root length of WT, *almt5almt9*, almt4almt9, almt4almt5 and almt4almt9almt5 lines grown *in vitro* on ½ MS media supplemented (A) or not (B) with 1% sucrose. The root length was measured using ImageJ 6 (n = 20), 9 (n = 20), 12 (n = 20) days after sowing. Data are mean + SD.

(H) *AtALMT4, ATALMT9 and AtALMT5* expression quantified by qRT-PCR. 40 days-old WT plants were grown in short days conditions (8h light). The ninth leaf of the rosette was collected at the end of the day (ED). 10 plants were collected and analysed pooled by two. Transcript accumulation was quantified after normalisation by the reference gene Yellow leaf specific gene (YLS8).

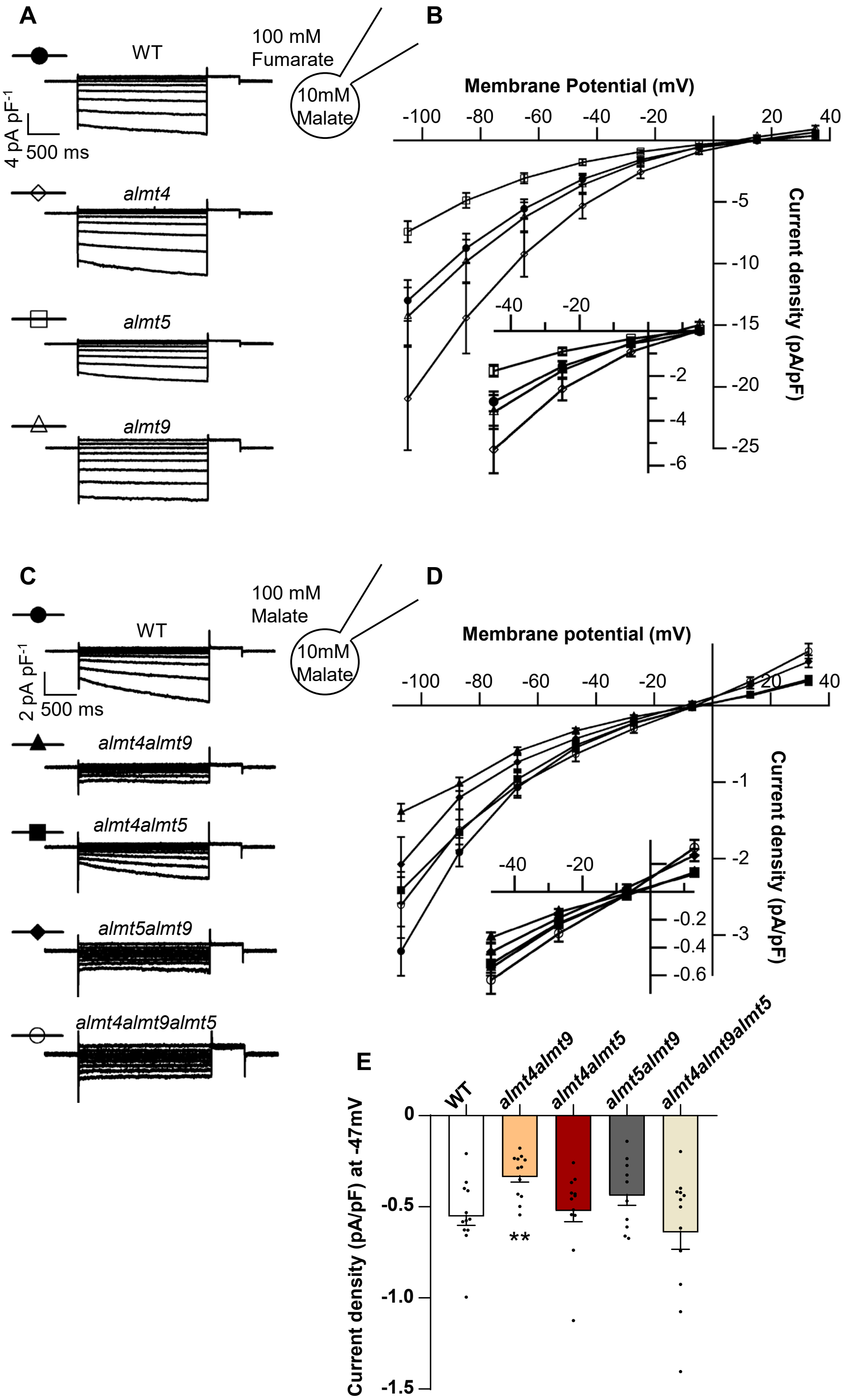

**Supporting Figure 4 : Vacuolar transport capacities of fumarate and malate by vacuolar *At*ALMT channels.**
(A and C), Representative whole-vacuole traces in cytosolic fumarate (A) or malate (C) conditions from WT, *almt4*, *almt5* *almt9*, *almt4almt9*, *almt4almt5*, *almt5almt9* and *almt4almt9almt5*. Starting at a holding potential set at 0 mV. 2 second voltage steps in -20 mV decrements were applied starting from +35 until -125 mV (A) or from +33 until -127 mV, each followed by a step at +35 mV (A) or (C) at -33 mV for 500 ms. (B and D) I/V curves of the mean current densities in cytosolic fumarate (B, WT: n=12; *almt4*: n=14; *almt5*: n=10; *almt9*: n=10) or cytosolic malate (D, WT: n=12; *almt4almt9*: n=12; *almt4almt5*: n=12; *almt5almt9*: n=10; *almt4almt9almt5*: n=12). (E) Mean current densities at -47 mV in cytosolic malate from D. (B, D and E) Each data is represented as follow mean ± SEM.

**Supporting Table 1** : Organic and inorganic anion contents mutants and in complemented lines.

| | Fumarate | Malate | Malate + Fumarate | Citrate | Chloride | Sulphate | Phosphate | Nitrate |
|---|---|---|---|---|---|---|---|---|
| WT | 10,4 ± 2,4 | 12,4 ± 3,2 | 22,8 + 4,0 | 14,1 ± 3,8 | 2,4 ± 0,7 | 10,7 ± 3,0 | 8,9 ± 2,5 | 168,1 ± 38,7 |
| *almt9* | 12,0 ± 2,2 | 11,2 ± 1,7 | 23,2 + 3,5 | 13,8 ± 2,1 | **1,9 ± 0,3 *** | 10,1 ± 1,5 | 8,0 ± 1,2 | 160,6 ±21,0 |
| *almt4* | 11,4 ± 3,8 | 11,6 ± 3,4 | 23,0 + 6,8 | 12,3 ± 3,1 | **1,8 ± 0,4 **** | 9,1 ± 2,8 | 7,3 ± 2,0 | 152,3 ± 40,0 |
| *almt5* | **4,3 ± 0,8 ***** | **16,2 ± 3,4 **** | 20,5 + 4,0 | 14,5 ± 3,2 | 2,3 ± 0,8 | 10,0 ± 2,0 | 8,1 ± 1,7 | 163,9 ± 31,9 |
| *almt5almt9* | **3,5 ± 0,6 ***** | 11,7 ± 1,7 | **15,3 + 2,2 **** | 15,4 ± 2,2 | **1,9 ± 0,3 *** | 13,0 ± 1,7 | 9,3 ± 1,2 | 155,2 ± 17,5 |
| *almt5almt9/ALMT9* | **4,8 ± 0,6 ***** | **15,9 ± 2,4 *** | 20,6 + 2,9 | 14,0 ± 1,7 | 2,0 ± 0,6 | 8,7 ± 1,7 | 7,5 ± 1,2 | 166,3 ± 20,3 |
| *almt5almt9/ALMT5* | 10,6 ± 3,3 | 11,7 ± 1,9 | 22,3 + 3,5 | 14,7 ± 1,6 | 2,4 ± 0,6 | 10,5 ± 1,9 | 7,7 ± 1,1 | 160,5 ± 15,5 |
| *almt4almt9* | 12,4 ± 2,9 | 10,6 ± 2,5 | 23,0 + 5,1 | 13,1 ± 2,5 | 2,1 ± 0,6 | 10,6 ± 2,5 | 8,1 ± 2,0 | 159,1 ± 33,4 |
| *almt4almt5* | **3,7 ± 0,7 ***** | **15,7 ± 2,5 *** | **19,5 + 3,1 *** | 14,4 ± 2,3 | 2,0 ± 0,5 | 10,1 ± 2,0 | 8,0 ± 1,5 | 169,9 ± 26,3 |
| *almt4almt9almt5* | **1,2 ± 0,2 ***** | **5,7 ± 1,7 ***** | **6,9 + 1,8 ***** | **7,5 ± 2,3 ***** | 2,4 ± 0,3 | **14,0 ± 3,6 *** | 7,3 ± 2,4 | 157,3 ± 28,0 |

Inorganic and organic anions content measurement of WT, *almt5, almt9, almt4, almt5almt9*, *almt4almt5, almt4almt9,* and complemented lines *almt5almt9/gALMT9* and *almt5almt9/gALMT5* 40 days-old rosettes grown in short days photoperiod (8h light). Entire rosettes were collected at the end of the day (ED) in the growth chamber. Each measurement correspond to n = 10 rosettes. Data were obtained by HPIC. Data are mean ± SD. Asterisks show a difference from the WT. Statistical analysis : Mann-Whitney test (*P<0.05;**P<0.01;***P<0.001) in comparaison to WT. The data of WT, *almt5*, *almt9*, *almt4*, *alm4almt5, almt4almt9*, *almt5almt9* and *almt4almt9almt5* in malate, fumarate, and malate + fumarate are the one show in Fig. 2D-F.

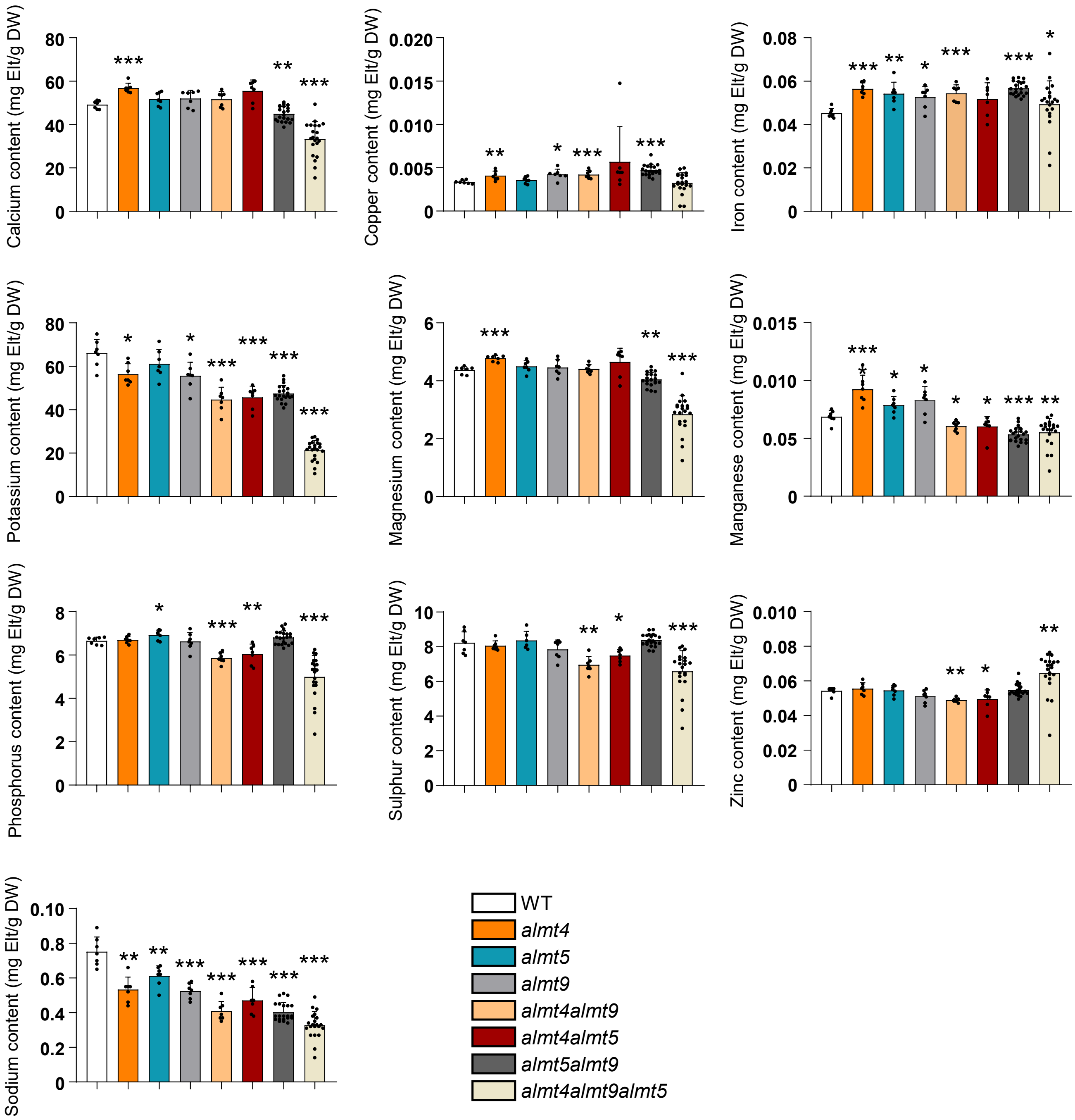

**Supporting Figure 5 : Cations contents in WT and ALMT mutants.**
Quantification of cations by ICP-OES in entire rosettes grown in short days conditions (8h light). Rosettes were collected at 40 days-old at the end of the day (ED). The data of potassium are visible Fig. 3. Asterisks show a difference from the WT. Statistical analysis : Mann-Whitney test (*P<0.05;**P<0.01;***P<0.001).

**Supporting Table 2** : Carbon (C) and Nitrogen (N) contents of complemented lines.

| | % C | δ13C | A%13C | % N |
|---|---|---|---|---|
| WT | 33,44 ± 0,92 | -31,35 ± 0,32 | 1,12 ± 0,00 | 6,35 ± 0,14 |
| *almt5* | 33,49 ± 1,18 | -31,41 ± 0,31 | 1,12 ± 0,00 | 6,26 ± 0,32 |
| *almt9* | 33,97 ± 0,89 | -31,41 ± 0,18 | 1,12 ± 0,00 | 6,26 ± 0,17 |
| *almt5almt9* | **34,76 ± 1,09 **** | **-29,86 ± 0,33 ***** | 1,12 ± 0,00 | **6,15 ± 0,19 *** |
| *almt5almt9/gALMT5 L9-6* | 34,03 ± 1,93 | **-31,6 ± 0,19 *** | 1,12 ± 0,00 | **6,01 ± 0,35 **** |
| *almt5almt9/gALMT9 L24-3* | 34,19 ± 0,63 | -31,59 ± 0,16 | 1,12 ± 0,00 | **6,20 ± 0,10 **** |

Carbon and Nitrogen content measurement of WT, *almt5, almt9*, *almt5almt9*, *almt5almt9/gALMT5 L9-6* and *almt5almt9/gALMT5 L24-3* 40 days-old rosettes grown in short days photoperiod (8h light). Entire rosettes were collected 5h after the beginning of the photoperiod in the growth chamber. Each measurement correspond to n = 10 rosettes. Data were obtained by mass spectrometry. Data are mean ± SD. Asterisks show a difference from the WT. Statistical analysis : Mann-Whitney test (*P<0.05;**P<0.01;***P<0.001). WT, *almt5*, *almt9* and *almt5almt9* data are the one show in Table 1.

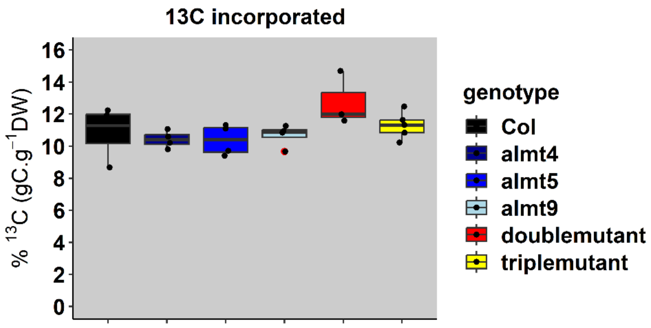

**Supporting Figure 6 : $^{13}$C allocation in *Arabidopsis thaliana* rosettes.**
A, Percentage of $^{13}$C incorporated in WT, *almt4*, *almt5*, *almt9*, *almt5almt9* and *almt4almt9almt5* lines after 4h of $^{13}CO_2$ incorporation. Plants were 6-week-old grown in short days (8h light). Plants were placed in a $^{13}CO^2$ chamber 2-3h after the beginning of the light period.

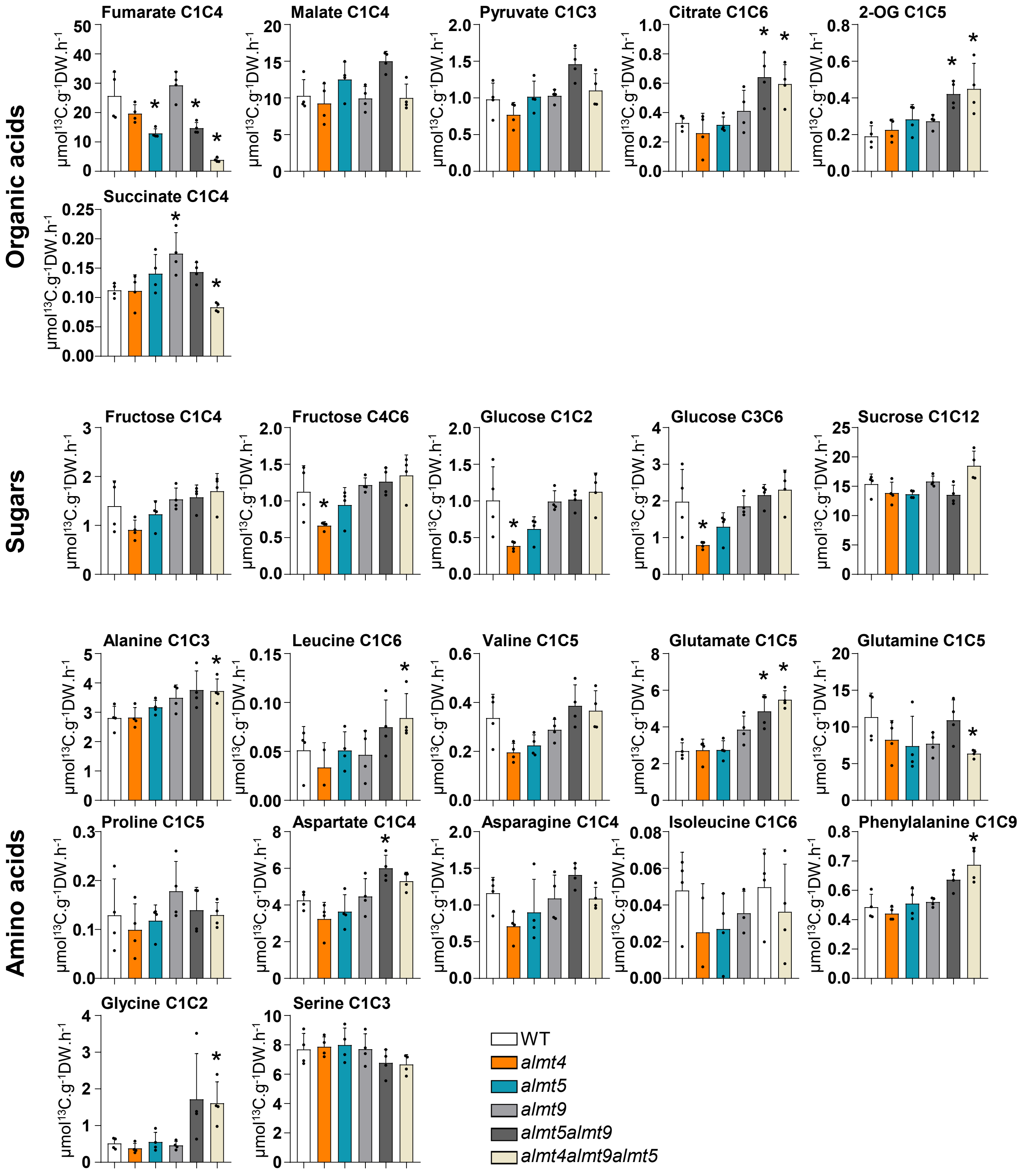

**Supporting figure 7 : $^{13}CO_2$ incorporation in *Arabidopsis thaliana* rosettes metabolism.**

$^{13}C$ incorporation quantification in metabolites and amino acids on WT, *almt4, almt5, almt9, almt5almt9* and *almt4almt9almt5* (n = 4 entire rosettes). Quantification was done by GC-MS after 4h of $^{13}CO_2$ incorporation. Statistical analysis: Mann-Whitney test (*P<0.05). Asterisks represents a significant difference compared to the WT. Results are expressed in µmol$^{13}$C.g$^{-1}$DW.h$^{-1}$.

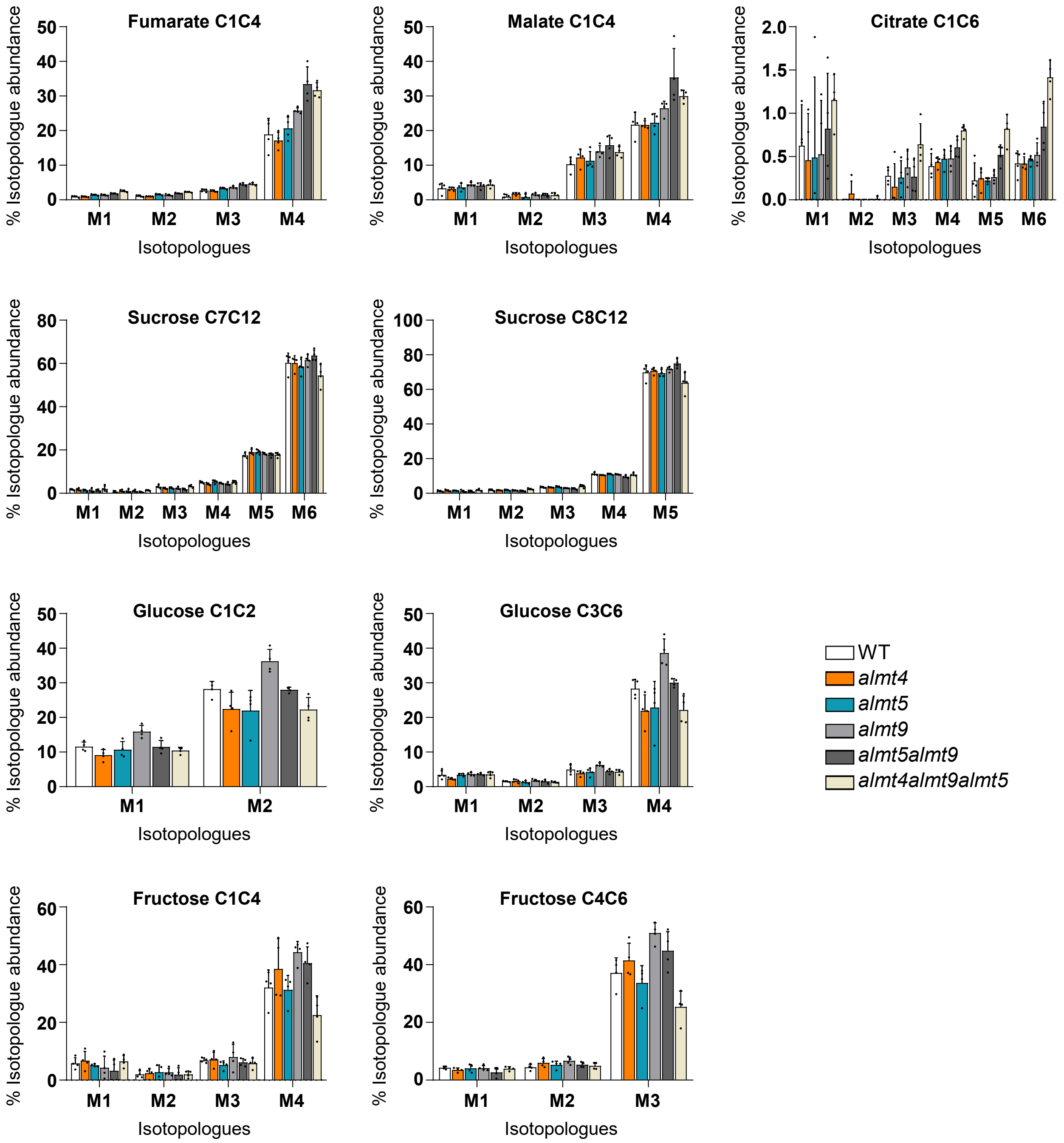

**Supporting figure 8 : Evaluation of carbon isotopologue distribution.**
Percentage of isotopologues abundance in WT, *almt4*, *almt5*, *almt9*, *almt5almt9* and *almt4almt9almt5* lines after 4h of $^{13}CO_2$ incubation. Plants were 6-week-old grown in short days (8h light). Plants were placed in a $^{13}CO_2$ chamber 2-3h after the beginning of the light period. Results are mean + SD of four independent biological replicates. The number of carbon of each metabolites is written on the title of the graphs.

**Supporting Table 3**: List of primers.

| AGI | Primers Name | Forward | Reverse | Purpose | Reference |
|---|---|---|---|---|---|
| At3g18440 | ALMT9-1 | ACCAGCCAAGAAGAGTCTCTGATG | AGTGAATGCACAATGCTTGAGAGC | qPCR | (Doireau et al., 2024) |
| At1g68600 | ALMT5-1 | AGAGATGTGGCCAAGTATGTCC | GAGACAACGAACTCGCACTTTC | qPCR | (Doireau et al., 2024) |
| At1g25480 | ALMT4-1 | ATCCTGGTTGGTGCTGGTATC | AGGTCTTCTCCAGCCCAAATG | qPCR | This publication |
| At5g47560 | tDT | CCGTCGAACACTACAACATCC | GCTGTTGTGGCGCAGATGC | qPCR | (Frei et al., 2018) |
| At5g50950.1 | FUM2 | CTGTGTTAGAGGCATTGAGGCC | GCATTGTCATAGCCAATTTTAGGA | qPCR | (Riewe et al., 2016) |
| At1g18420.1 | ALMT3 | GGCTTATCCTACAGAGCAGAGGCT | TCAGAGCCAAACCCATCTTC | qPCR | (Eisenach et al.,2017) |
| At5g08290 | YLS8 | AAGAGCGTCTCGTCGTCATTCG | ACGCAAGCACCTCATCCATCTG | qPCR | This publication |
| At2g17470.2 | ALMT6 | CCGTTGCATGATGCTAGTAAATAC | TGATGATGGTTTGCTCGAAA | qPCR | (Eisenach et al.,2017) |
| | almt5-1_fwd / almt5_1463_rev | ATGGGAGGTAAAATGGGATCA | CGGATCTGTACCCGCTGTAT | Genotyping by PCR | |
| | Wisc_p745 / almt5_1463_rev | AACGTCCGCAATGTGTTATTAAGTTGTC | CGGATCTGTACCCGCTGTAT | Genotyping by PCR | |
| | almt9_959_fwd / almt9_2446_rev | CCCGTCGATGAAAGCTTATG | CATCCCAAAACACCTACGAATCTT | Genotyping by PCR | |
| | almt9_959_fwd / LBb1.3 | CCCGTCGATGAAAGCTTATG | ATTTTGCCGATTTCGGAAC | Genotyping by PCR | |
| | almt4_394_fwd / almt4_1013_rev | TTCGATCAAATCTTCCGGATTT | CTTCAAGTGAATTGGCCACAC | Genotyping by PCR | |
| | almt4_394_fwd / LBb1.3 | TTCGATCAAATCTTCCGGATTT | ATTTTGCCGATTTCGGAAC | Genotyping by PCR | |
